# Purifying selection against spurious splicing signals contributes to the base composition evolution of the polypyrimidine tract

**DOI:** 10.1101/2022.01.29.478184

**Authors:** Buŗcin Yıldırım, Claus Vogl

## Abstract

Among eukaryotes, the major spliceosomal pathway is highly conserved. While long introns may contain additional regulatory sequences, the ones in short introns seem to be nearly exclusively related to splicing. Although these regulatory sequences involved in splicing are well-characterized, little is known about their evolution. At the 3’ end of introns, the splice signal nearly universally contains the dimer *AG*, which consists of purines, and the polypyrimidine tract upstream of this 3’ splice signal is characterized by over-representation of pyrimidines. If the over-representation of pyrimidines in the polypyrimidine tract is also due to avoidance of a premature splicing signal, we hypothesize that *AG* should be the most under-represented dimer. Through the use of DNA-strand asymmetry patterns, we confirm this prediction in fruit flies of the genus *Drosophila* and by comparing the asymmetry patterns to a presumably neutrally evolving region, we quantify the selection strength acting on each motif. Moreover, our inference and simulation method revealed that the best explanation for the base composition evolution of the polypyrimidine tract is the joint action of purifying selection against a spurious 3’ splice signal and the selection for pyrimidines . Patterns of asymmetry in other eukaryotes indicate that avoidance of premature splicing similarly affects the nucleotide composition in their polypyrimidine tracts.

## Introduction

Noncoding sequences, both intergenic and intronic, comprise a big proportion of eukaryotic genomes. Some noncoding sequences influence the centrally important processes of chromosome assembly, DNA replication, and gene expression (Ludwig, 2002; Pennacchio & Rubin, 2001). Other noncoding sequences were identified as nearly unconstrained and have been used as a neutral reference for inference of demography and selection in many population genetic analyses (*e.g.,* Parsch et al., 2010; Lawrie & Petrov, 2014). Introns are known to contain conserved *cis*-regulatory motifs that are necessary for splicing. Almost all eukaryotes contain a mixture of long and short introns . It is thought that the recognition and removal of introns during pre-mRNA splicing differs between long and short introns. In long introns, exons define the recognition unit (“exon definition”): the 5’ end of the exon (which corresponds to the 3’ end of the previous intron) and the 3’ end of the exon (which corresponds to the 5’ end of the next intron) are recognized as a pair (Berget, 1995). In short introns, introns define the recognition unit (“intron definition”): 5’ and 3’ splice sites of the same intron are recognized as a pair (Talerico & Berget, 1994). This implies that intron length will influence the selection pressure on splicing motifs: with intron definition, *i.e.,* in short introns, the splicing information resides mainly in the intron. Stronger splicing-coupled selection in short introns of several eukaryotic species, including *Drosophila melanogaster* compared to long introns has been documented (Farlow et al., 2012). Furthermore, overall selective constraint increases with the length of introns, suggesting that the number of functional elements not related to splicing is increased (Haddrill et al., 2005; Belshaw & Bensasson, 2006).

Regardless of their length, all introns contain common conserved sequences necessary for splicing (Green, 1986). These include the 5’ (or donor) splice site, the 3’ (or acceptor) splice site and the branch point. The 5’ splice site has a consensus sequence of *G|GTRAG*; the 3’ splice site a consensus sequence of *Y AG|G* (where *|* denotes the exon-intron or intron-exon boundary, respectively, and *Y* defines a pyrimidine, *i.e.,* one of the bases *C* or *T*, and *R* a purine, *i.e.,* one of the bases *A* or *G*) (Breathnach & Chambon, 1981; Mount, 1982). Additionally, there is a pyrimidine-rich region of variable length, upstream of the 3’ splice site, referred to as the polypyrimidine tract (abbreviated as 3PT in this article). The pre-mRNA splicing pathway has been shown to involve two main reactions which are universal to eukaryotic organisms (Ruskin et al., 1984; Green, 1986) (fig. 1). In the first reaction, the 5’ splice site is cleaved and the RNA forms a loop (lariat) by attaching bluntly, *i.e.,* without base pairing, to the branch point. (We will refer to this region as 5’ loop region, abbreviated as 5LR in this article.) Thereafter, splicing occurs at the 3’ end, and exons are ligated (Green, 1986). Experiments characterizing the splicing intermediates have shown that the order of the splice-site cleavage is highly conserved: no 3’ splice-site cleavage is observed without the cleavage of the 5’ splice site (Padgett et al., 1984; Grabowski et al., 1984; Ruskin et al., 1984).

**Figure 1:**
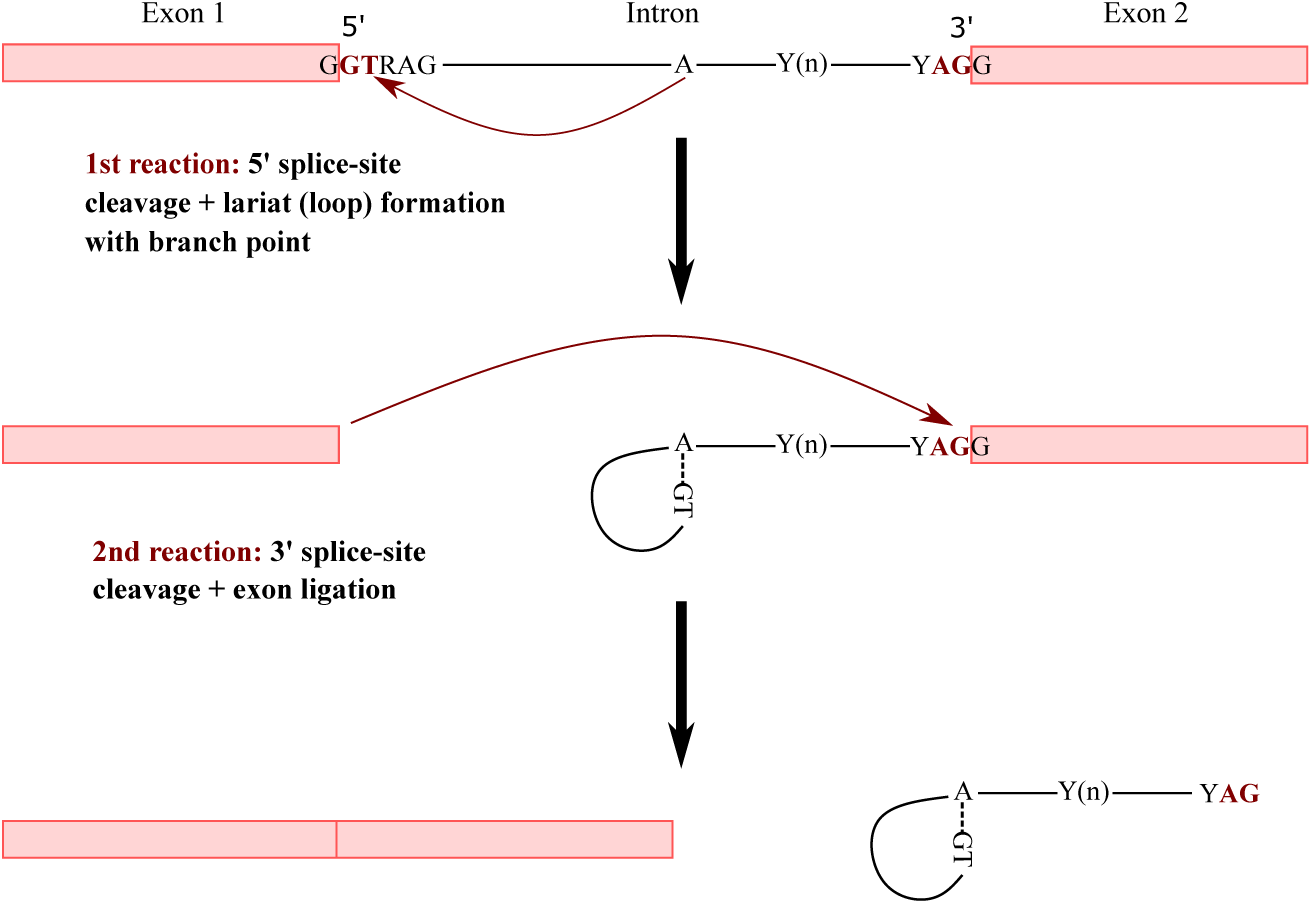
Representation of the pre-mRNA splicing pathway (adapted from Green, 1986)

The relative importance of the *cis*-regulatory motifs in introns has been elucidated by mu-tagenesis studies (for reviews see Green, 1986; Padgett et al., 1986). Even though both ends of the intron are marked with long consensus sequences required for maximal splicing efficiency, the *GT* and *AG* dinucleotides immediately adjacent to the 5’ exon-intron and 3’ intron-exon boundary, respectively, seem to be the most functionally important and conserved motifs (Breathnach & Chambon, 1981). Mutations in these nearly invariant dinucleotides completely inactivate the authentic splice sites and often result in the activation of cryptic (alternative) splice sites, while mutations at other positions within these signals have lesser effects (Green, 1986; Padgett et al., 1986). The sequence of the branch point is not conserved universally among eukaryotes: Generally an adenosine residue is surrounded by rather variable sequence elements (Ruskin et al., 1985).

While the sequence of the branch point may be variable, the minimum distance to both the 5’ and 3’ ends seems to be strongly constrained. Decreasing the distance between the 5’ splice site and branch point prevents accurate splicing or leads to the activation of an upstream cryptic 5’ splice site (Green, 1986). While the total length of the 5LR seems constrained, its base composition seems to evolve neutrally. Similarly, the position of the branch point relative to the 3’ end, and thus the length of the 3PT, is conserved (Ruskin et al., 1985). In contrast to the 5’ side, not only the length, but also the base composition of the 3PT seem important, as some mutations in the polypyrimidine tract and the decrease in its length reduce splicing efficiency (Ruskin & Green, 1985; Coolidge et al., 1997), while increasing the number of consecutive pyrimidines enhances the removal of the intron (Guo et al., 1993; Coolidge et al., 1997). Several trans-splicing factors preferentially bind to the 3PT by identifying motifs, including the essential pre-mRNA splicing factor U2AF65 which is a part of the U2AF heterodimer (Zamore et al., 1992). The preference for specific sequences of these proteins is thought to explain the constraint on the base composition of 3PT.

Experimental studies revealed that the *AG* dimer downstream of the branch point is found by a 5’-to-3’ scanning mechanism and the base preceding this downstream *AG* creates competition (Smith et al., 1993). Recognition of the *AG* splicing signal fails if it is too close to the branch point, explaining the constraint on the length of this region. Notably, decreasing the pyrimidine content from 70% to 30% in the polypyrimidine tract did not affect the recognition of *AG* and the 3’ splice-site cleavage (Smith et al., 1989, 1993). Other mutagenesis studies also reported similar results: the introduction of purines into the polypyrimidine tract is only detrimental if it changes the length of 3PT drastically and if the number of consecutive uridine bases decreases (Roscigno et al., 1993; Coolidge et al., 1997). This suggests high variation in the base composition of the 3PT, which is supported by studies showing diversity in nucleotide composition of polypyrimidine tracts among introns within a species (Coolidge et al., 1997; Green, 1991) and among eukaryotic species (Sickmier et al., 2006; Schwartz et al., 2008). At first, this seems at odds with what is known about the trans-acting elements associated with the 3PT, which recognize motifs in this region. Yet, there are multiple polypyrimidine tract binding proteins and each with a different binding affinity to different nucleotide compositions (Singh et al., 1995, 2000). Moreover, even though the U2AF65 protein has a preference for uridine-rich sequences, it has been shown to tolerate diverse 3PT nucleotide sequences with different affinities (Zamore et al., 1992; Green, 1991). Sickmier et al. (2006) showed that U2AF65 recognizes uridines through hydrogen bond interaction, rather than selecting the smaller shape of pyrimidines compared to purines. These findings might explain its flexibility in recognizing different 3PT sequences.

All of these results suggest that the base composition of the 3PT is not just a result of selection for pyrimidines. While generally preferring pyrimidines and especially uridine-rich sequences, trans-acting elements associated with the 3PT exert relatively weak selective pressure on 3PT sequence evolution. On the other hand, the scanning mechanism downstream of the branch point to identify the conserved *AG* dinucleotide creates a strong selective pressure to avoid a premature 3’ splice site. Considering that the *AG* dimer consists of only purine bases, avoidance of this dimer would also affect the base composition in favor of pyrimidines. Consequently, there might be three possibilities for the evolution of base composition in the 3PT: I) selection for *Y* s fully explains the pattern; II) avoidance of the canonical 3’ splice site motif *AG* fully explains the over-representation of *Y* s; III) both selection for *Y* s and selection against *AG* are necessary to explain the pattern in the 3PT.

Obviously, un-spliced or mis-spliced introns may be deleterious for the cell and organism (Jaillon et al., 2008), so that selection should maintain efficient splicing, which requires the *cis*-regulatory motifs and binding of the splicing machinery to this specific RNA sequence motifs. The nucleotide composition of the region of the 3PT should show evidence of this splicing-coupled selection for and against specific motifs. Several studies have identified splicing-related sequence motifs using DNA-strand asymmetry patterns (Zhang et al., 2008; Farlow et al., 2012). According to Chargaff’s second parity rule, mono- or oligonucleotides under neutral evolution should have the same frequency as their reverse complement (Mitchell & Bridge, 2006). In many organisms, deviations from Chargaff’s second parity rule are observed, associated with either selection-driven or neutral processes. Neutral processes, *e.g.,* replication or transcription-coupled asymmetries, are attributed to different mutation and repair pressures between lagging and leading strands for replication or coding and non-coding strands for transcription (Touchon et al., 2003; Green et al., 2003). If deviations are mainly driven by transcription-coupled repair, one would expect a constant asymmetry score along the transcribed region and no over- or under-representation of motifs. On the other hand, deviation from Chargaff’s second parity rule due to positive or negative selection leads to over- or under-representation of the selected motif. A comparison of counts of motifs to counts of their reverse complements may identify elements involved in splicing.

While investigating strand asymmetry patterns might reveal splicing-associated motifs, estimating the strength of selection would require a reference sequence class free from selective pressure. As mentioned, the loop formation between the 5’ splice site and branch point does not involve base pairing. Thus selection on nucleotide composition due to splicing in the 5LR should be minimal between the 5’ splice signal and the branch point, as long as a proper length is maintained. Additionally, in *Drosophila*, studies showed that sites at positions 8-30 of introns shorter than 65 bp have a higher divergence and polymorphism compared to other regions in introns (Halligan & Keightley, 2006; Parsch et al., 2010; Clemente & Vogl, 2012). This has been interpreted as evidence for little or no selection. In these short introns, a ratio of about *AT* : *GC* = 2 : 1 likely reflects mutation bias (Haddrill et al., 2005; Clemente & Vogl, 2012), while the *GC* content increases with increasing intron length likely due to selection (Haddrill et al., 2005). Lawrie and colleagues (Lawrie et al., 2013; Machado et al., 2020) later utilized these short intronic regions to infer directional selection on constrained sites in the presence of mutation bias in *Drosophila*, showing the utility of these intronic sites as a reference for inferring weak selection (Vogl & Bergman, 2015). It seems that the base composition and higher-order oligonucleotide motifs in the 5LR of short introns of *Drosophila* can be used as a neutral reference. Recent studies showed that *Drosophila* performs transcription-coupled repair (Deger et al., 2019; Törmä et al., 2020) and this might also lead to deviation from strand symmetry in the 5LR (Bergman et al., 2017). Yet it can be assumed that this effect would be similar along the intron, and as long as selection is absent in the 5’ region, it could be used to control and quantify the effects of splicing-coupled selection in other parts of the intron.

In this study, we investigate the evolution of base composition in the 3PT and aim to distinguish among three hypotheses proposed to explain its evolution. We begin by using strand asymmetry patterns to assess whether selection acts on specific oligonucleotide motifs, not just on pyrimidine monomers. Specifically, we compare complementary motifs within the 3PT and reveal the effect of splicing-coupled selection acting on splice signal-associated motifs, thus rejecting hypothesis I, which posits selection only on monomers. Furthermore, we quantify the strength of selection acting on each motif in *Drosophila* by comparing asymmetry patterns of motifs between the 3PT and the presumably neutral 5LR. Lastly, to further differentiate among hypotheses, we use inferred selection coefficients modeled through fixation probabilities to compare three hypotheses and perform simulations under these models. Our findings suggest that avoidance of the *AG* dimer, along with selection on the monomer level, is necessary to explain the base composition of the 3PT.

## Materials and Methods

### Data used in the analyses

We analyzed whole genome data of a Zambian *D. melanogaster* population (Lack et al., 2015) and *D. simulans* populations from Madagascar (Rogers et al., 2014; Jackson et al., 2017). For *D. melanogaster*, the dataset consists of 197 individuals for autosomes and 196 for the X chromosome. For analyses requiring comparison between chromosomes, the individual missing from the X chromosome data was also excluded from the autosomal data. Additionally, we repeated the analyses for 69 Zambian individuals that showed no evidence of admixture with European populations according to Lack et al. (2015). The *D. simulans* dataset includes 21 individuals for each chromosome. Sequences have been obtained as consensus FASTA files. Using annotations from the reference genomes of *D. melanogaster* (r5.57 from http://www.flybase.org/) and *D. simulans* (Hu et al., 2013), intron coordinates were extracted and alignments of all samples were created. Due to alternatively spliced isoforms in the GFF file (see https://www.ensembl.org/info/website/upload/gff.html; last accessed November 1, 2020), multiple entries may exist for the same intron. To avoid including the same intron sequence more than once, only one entry of an intron with the same coordinates was used. If the annotation information was coming from the non-coding strand, the sequence was reverse-complemented, so that the direction of all alignments of transcripts is from 5’-to-3’. The results presented in the main text focus on the *D. melanogaster* dataset, while the *D. simulans* dataset is used for comparison and can be found in the supplement.

In *Drosophila* the length distribution of introns seems to fall into a majority of short and a minority of long introns, but the boundary between these two classes is unclear. Therefore we tried to define an upper length limit for the short intron class by binning according to length and then checking the nucleotide composition. For this analysis, we also included fifty bases from the preceding and following exons.

We differentiated i) a presumably neutral region between the 5’ end and the branch point, the 5LR and ii) a presumably selected region through the 3’ end with an excess of pyrimidines, the 3PT. The 5LR is characterized by a ratio of approximately *AT* : *GC* = 2 : 1, which possibly reflects the mutation bias; the 3LR by a high pyrimidine content, which possibly reflects selection. Note that the branch point cannot easily be defined by a characteristic sequence pattern. To compare regions among introns of varying length, consensus positions from the 5’ and 3’ splice sites were defined among all length classes. Additionally, an eight-bp long region that symmetrically straddled the 3’ junction (4 bp into the intron and 4 bp into the exon) was extracted.

Sequences were filtered out if they overlapped with annotated coding sequences or if they contained undefined nucleotides (*N*) in at least one of the individuals in the alignment. Furthermore, only alignments with full-length stretches (23 bp for the 5LR and 10 bp for the 3PT; see Results) were used for further analyses. Following these filtering steps, position-weight matrices and consensus sequences were created for the remaining alignments, resulting in 14,762 and 2,036 sequences for autosomal and X-linked 3PTs, respectively, and 11,938 and 1,556 sequences for autosomal 5LRs, respectively. By scanning these consensus sequences, all possible dimer (4^2^ = 16), trimer (4^3^ = 64), and tetramer sequences (4^4^ = 256) were counted, separately for each region and chromosome. The expected proportion of oligomers was calculated from the base composition of the particular region. Results from autosomal introns are presented in detail in the text; results from X-linked introns in the supplement.

To evaluate whether alternative splicing influences our results, we binned introns that are in phase, *i.e.,* of length 3*n* + 0 (phase 0), and introns out of phase, *i.e.,* of length 3*n* + 1 or 3*n* + 2 (non-phase 0). We then repeated analyses for datasets including i) only non-phase 0 introns in *D. melanogaster* ii) only phase 0 introns in *D. melanogaster* and excluding iii) common phase 0 introns between *D. melanogaster* and *D. simulans*, and compared the results to the full dataset.

### Tests of strand symmetric evolution in the 5LR

Under neutrality and without transcription-associated mutation bias, counts of a motif and its reverse are expected to be identical. We examined this expectation in the 5LR with two, not mutually exclusive, tests:

- Test 1: a chi-square test to see whether forward and reverse oligonucleotide sequences are equally represented.
- Equivalence test: a test proposed by Afreixo et al. (2013), to see whether there are significant deviations from a 1:1 ratio between forward and reverse sequences. The procedure consists of obtaining confidence intervals for ratios (Katz et al., 1978) and checking if they are contained in the tolerance range, (1*/δ, δ*), where *δ* represents a small tolerance to conclude equivalence for a ratio. If so, equivalence can be assumed.

The lower and upper values of the confidence intervals for the ratio of motifs, for a given *z*, are calculated as:

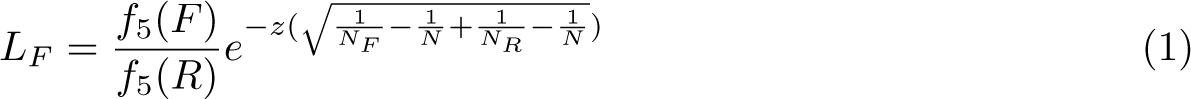

and

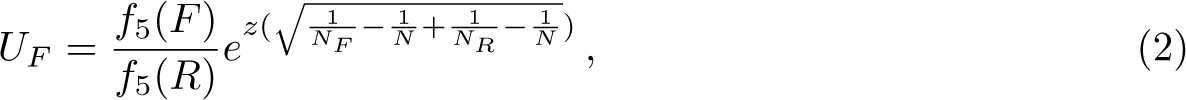

where *f*_5_(*F*)*/f*_5_(*R*) is the ratio between two proportions of forward and reverse sequences from 5LR, *N_F_* and *N_R_* is the frequency of the forward and reverse sequences, respectively and *N* is the total number of oligonucleotide occurrences. To assume practical equivalence, we used a stringent *δ* = 1.1 (Thanassoulis & Vasan, 2010).

### Controlling for GC-biased gene conversion in the 5LR

A consequence of the repair of double strand breaks is gene conversion. Heteroduplex mismatches formed during the repair of double strand breaks can involve either pairing between the bases *G* and *C* (strong: S:S), *A* and *T* (weak: W:W) or between strong and weak bases, S:W. Preferential resolution of the S:W mismatches into *G* : *C* rather than *A* : *T* leads to *GC*-biased gene conversion (gBGC) (Marais, 2003). Since, gBGC only affects S:W mismatches they are referred to as *GC*-changing, while the others (S:S, W:W) are called *GC*-conservative. This categorization allows us to use unpolarized data to estimate *GC* bias while considering the site frequency spectra (SFS) of all six possible nucleotide pairs (Borges et al., 2019). The efficiency of gBGC (quantified by *B*) is directly related to effective population size (*N_e_*), thus parametrized by the product of *N_e_*and conversion bias (*b*). To infer the strength of gBGC in the 5LR, we used the maximum likelihood estimator of Vogl & Bergman (2015) and applied it to the SFS from *GC*-changing mutations of the 5LR. The SFS from *GC*-conservative mutations was considered as putatively neutral control (*B* = 0). We performed likelihood-ratio tests (LRT) to compare between the different nested models.

### Calculation of asymmetry scores

Strand asymmetry of mono- and oligonucleotides for each region was calculated as

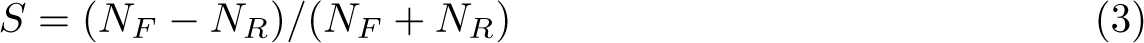

Mononucleotide asymmetries *T vs. A* and *C vs. G* are denoted as *S_T A_*and *S_CG_*, respectively (Touchon et al., 2004). Additionally, at the mononucleotide level, we also calculated asymmetry scores for each position in the 5LR, the 3PT, and the 3’ junction and documented the change in the scores with position via regression analysis to study whether there is a dependence on distance to the splice site. The expected strand asymmetry of an oligonucleotide was predicted from the base composition of the region under consideration.

We also obtained the short introns of six additional eukaryotes (human, sea urchin, *Caenorhabditis elegans*, moss, rice, and *Arabidopsis thaliana*) from the exon-intron database, EID (Shepelev & Fedorov, 2006) to document the strand asymmetry patterns in polypyrimidine regions of other species. The length range for the short intron class was defined separately for each species depending on the length distribution of introns and the nucleotide composition similar to that described for *Drosophila*. Inside these introns, probable polypyrimidine tracts were again characterized by a high pyrimidine content, and asymmetry scores of oligonucleotide motifs with lengths two and three were calculated. The number of introns (3PT region) used for each species; Human: 12740, Sea Urchin: 21285, Rice: 24137, *Arabidopsis*: 55354, Moss: 50585, *C. elegans*: 54082. Additionally, we obtained the introns of two yeast species, namely *Saccharomyces cerevisiae* and *Lachancea thermotolerans*. The genomic sequence and annotations for *S. cerevisiae* were extracted from the *Saccharomyces* genome database (strain S288C, https://www.yeastgenome.org/, last accessed January 31, 2023). For *L. thermotolerans*, the genomic sequence were retrieved from YGOB (http://ygob.ucd.ie/, version 7, last accessed January 31, 2023), and annotations were obtained from Hooks et al. (2014).

Compared to other eukaryotes yeast genomes contain very few introns (*∼* 300). Therefore we used all the annotated and fully sequenced introns for these species and extracted the pyrimidine-rich region at the 3’ end (277 and 216 introns for *S. cerevisiae* and *L. thermotolerans*, respectively).

### Quantification of selection strengths via strand asymmetry

Motifs distinguished from others by their strand asymmetry in the previous step can be functionally constrained due to splicing-coupled selection, which would lead to either under- or over-representation of pairs of symmetric sequences. We quantified the selection strength causing this asymmetry on particular motifs (denoted as *M*) using the 5LR as background:

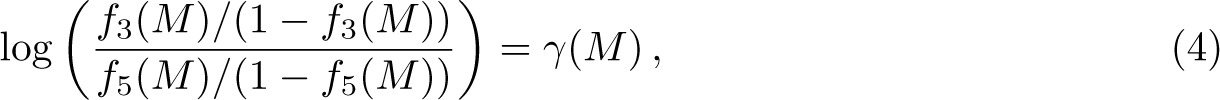

where *f*_3_(*M*) and *f*_5_(*M*) represent the proportion of motifs from 3PT and 5LR, respectively, and *γ* = 4*N_e_s* is the scaled selection strength, *i.e.,* the per generation selection strength *s* scaled by the effective population size *N_e_*. The method does not require complete strand symmetry in the 5LR, but instead assesses the selective force leading to either depletion or excess of the particular motif in the 3PT compared to the 5LR, in the presence of non-selective forces such as mutation bias.

To create confidence intervals (CIs) around the estimates, we bootstrapped by sampling introns (both 5LR and 3PT datasets) with replacement 1000 times. The size of the each bootstrap sample was approximately same with the original datasets. Selection coefficients were recalculated for each motif from these bootstrap samples, the 2.5% and 97.5% quantiles of the resulting distributions were used as lower and upper bounds of 95% CIs.

### Hypothesis testing via population genetic modeling

On top of the strand asymmetry patterns to detect and quantify the selection on motifs, we also applied a method to differentiate the relative contribution of selection on monomers and dimers to the base composition of 3PT. We assumed that mutation rates are low, such that the segregation of more than two alleles in the population is negligible. Selection was modeled through its effect on fixation probabilities (Kimura, 1962); with relatively small selection the fixation rate is:

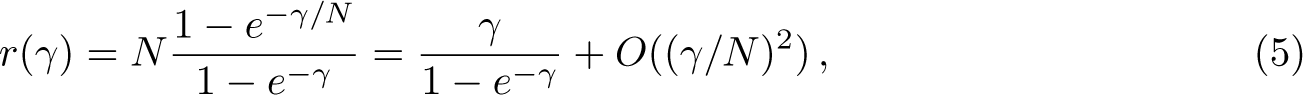

Under neutrality, the mutation rate matrix is defined as:

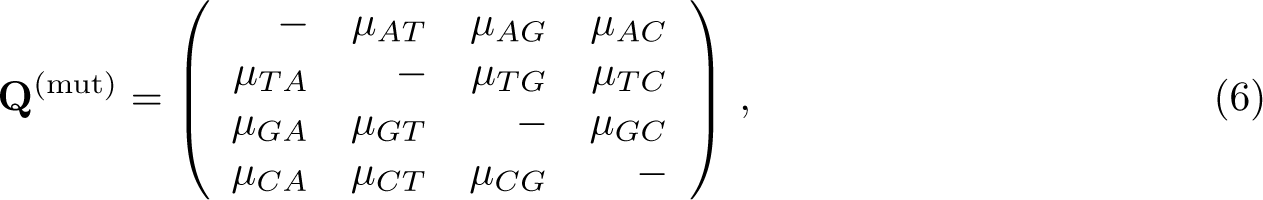

where rows and columns are ordered as (*A, T, G, C*) and diagonal elements correspond to minus the sum of the other elements. Next, we considered selection on monomers. We modelled selection effects among the four bases as a vector with four elements, which sum to zero: (*γ_A_, γ_T_, γ_G_, γ_C_*) with *_i_ γ_i_* = 0. We obtained the mutation-selection rate matrix by replacing *µ_ij_* with *r*(*−γ_i_* + *γ_j_*)*µ_ij_*:

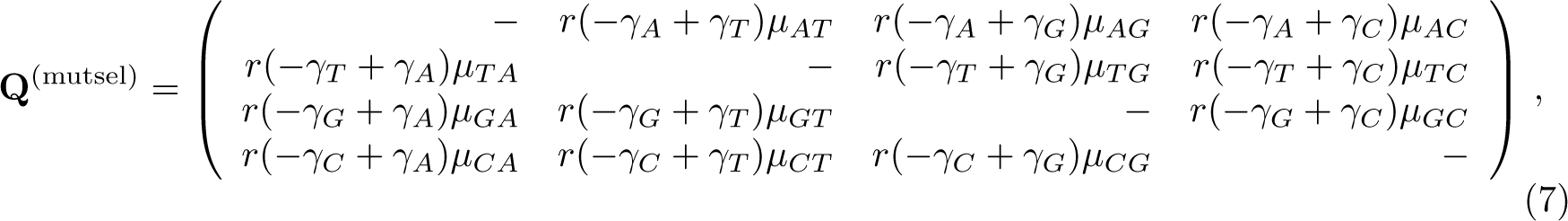

In addition to the monomer effects, the dimer *AG* was assumed to be selected with strength *γ_AG_*. Selection on monomers is local, while selection on dimers is determined by the sequence context. Conditional on an *A* preceding the focal position, we modified the column *µ_iG_* to *r*(*−γ_i_* + *γ_G_* + *γ_AG_*)*µ_iG_* and the row *µ_Gj_* to *r*(*−γ_G_* + *γ_j_ − γ_AG_*)*µ_Gj_*. Similarly, conditional on a *G* following the focal position, we modified the column *µ_iA_* to *r*(*−γ_i_* + *γ_A_* + *γ_AG_*)*µ_iA_* and the row *µ_Aj_* to *r*(*−γ_A_* + *γ_j_ − γ_AG_*)*µ_Aj_*. As above, the entry on the main diagonal balances the rest of the entries in the row.

We assumed a strand symmetric mutation rate matrix, where *A* and *T* or *G* and *C* nucleotides can interchange (*i.e., µ_AT_* = *µ_T A_*) and inferred this matrix from the 5LR of short introns by using the ML estimators from Vogl et al. (2020). We also obtained the joint frequency matrix of the four bases at position *i* and *i* + 1 (1 *≤ i ≤* 9) for each position in the 3PT. Given these we inferred the mutation selection matrix for each position with selection coefficients maximizing the likelihood calculated from the joint frequency and transition matrices of four bases under selection. We performed the inference under three different models, corresponding to three hypotheses, and compared the likelihoods via likelihood ratio tests. The first hypothesis considered only selection on monomers, the second only on the *AG* dimer, the third selection on both monomers and the *AG* dimer. To create confidence intervals around the parameter estimates of the best fitting model, we bootstrapped by sampling introns with replacement 1000 times.

With the inferred values, we simulated a stretch of DNA of length *L* = 10 indexed by *l*. To ensure that all positions are equal, the stretch was joined circularly, such that the next position after *l* = 10 was *l* = 1. We initialized each position *l* by sampling a base from the stationary distribution of the mutation matrix **Q**^(mut)^. Then we iterated by calculating an *L ×* 4 matrix consisting of a vector of mutation-selection rates for each position conditional on its state and the neighboring states, by first calculating the vector of mutation rates away from the current base, while setting the probability of remaining at the current base to zero, and then multiplying these probabilities by the fixation probabilities conditional on the neighboring states. The rate until the next change corresponds to the sum of this matrix; the position *l* was sampled from the relative rates of change at the *L* positions; and the new base corresponds to the relative probability of the three possible newly mutated bases at that position. This sequence was iterated 10^5^ times. After a burn-in period of 10^3^ iterations, we calculated the base composition at each position, specifically the joint frequency distributions at position *i* and *i* + 1 for the parameters estimated for each hypothesis, and compared them with the empirical data.

## Results

### Region definition inside short introns

The distribution of intron lengths in *D. melanogaster* has a very high mode around minimal lengths, creating a “short intron” class (Mount et al., 1992) and a long tail of longer introns. While the lower limit of the short intron class is given by the data, the upper limit is defined differently in studies of intron evolution (*e.g.,* Halligan & Keightley, 2006; Lawrie et al., 2013). Nevertheless all studies agree on patterns specific for the presumably neutral part of short introns: while the first seven bases at the 5’ end of each intron are affected by the presence of splice sites, thereafter mutation pressure in short introns leads to an *AT* -rich base composition of approximately *AT* : *GC* = 2 : 1 (Haddrill et al., 2005; Parsch et al., 2010). This matches our findings: bases in positions 8-30 of introns shorter than 65 bp exhibit a consistent *AT* : *GC* ratio of approximately 2 : 1. For introns with lengths between 65 to 85 bp, this presumably neutral region stretches out until position 40. In longer introns of more than 75 bp length, the base composition in positions 8-30 approaches a ratio of *AT* : *GC* = 1 : 1 (fig. 2A), presumably reflecting the effect of selection. We therefore used the ratio of *AT* : *GC* = 2 : 1 in positions 8-30 to differentiate short from long introns and included only short introns up to 75 bp in further analyses and extracted sequences between positions 8-30 as a proxy of a neutrally evolving region (fig. 3).

**Figure 2:**
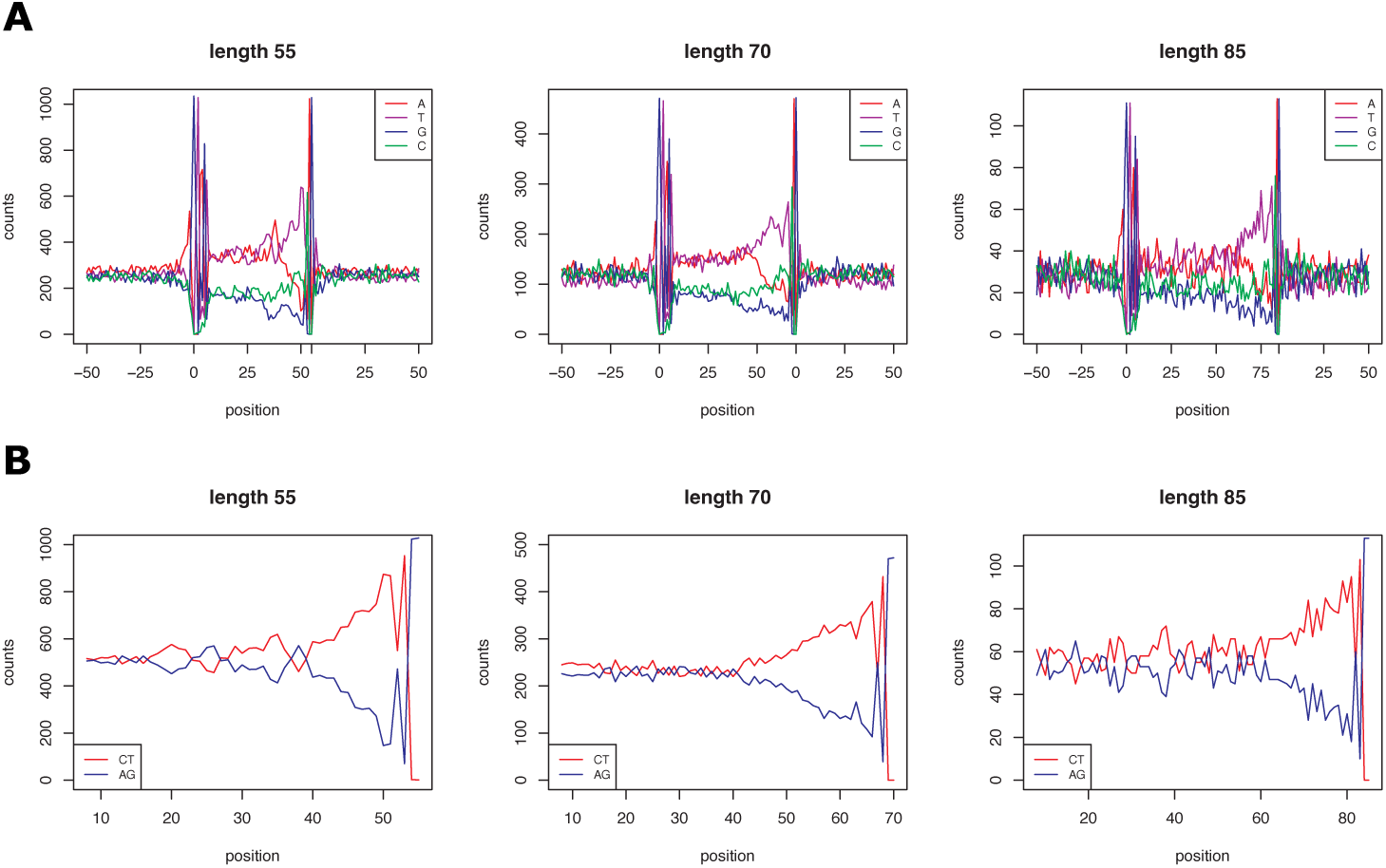
(A) Nucleotide composition around exon-intron junctions. Positions depicted as 0 correspond to the junctions between intron and exons. (B) *AG* (purines, blue lines) and *CT* (pyrimidines, red lines) content per position. Only one candidate length class were chosen to visualize the patterns with increasing length (55 bp, 70 bp, 85 bp). Total numbers of sequences used for each length class are 1044, 474 and 117, respectively

**Figure 3:**
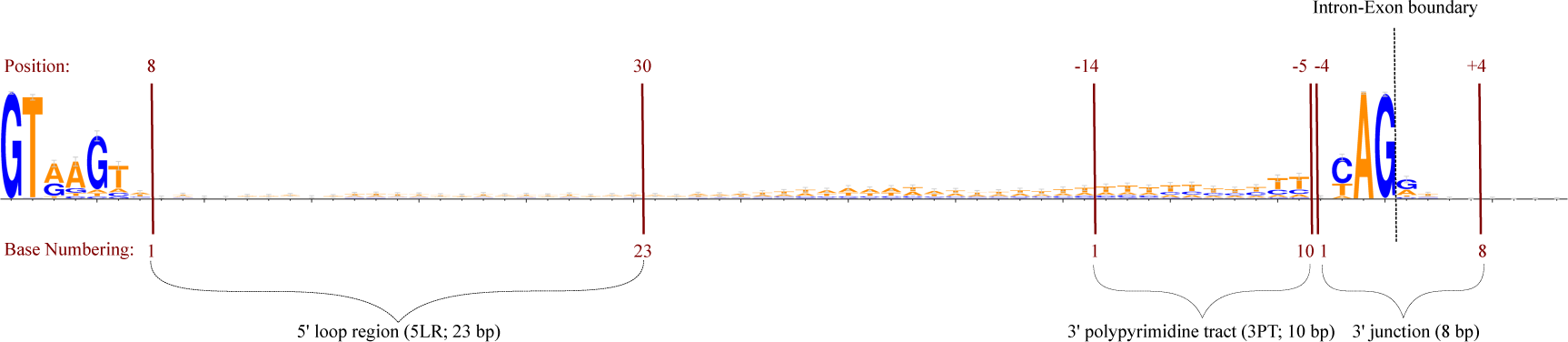
Representation of the regions analyzed. The base numbering of each region is ascending as going from 5’ to 3’.

We next searched for a region possibly under splicing-coupled selection inside short introns. The proportion of purine bases (*A* and *G*) and pyrimidine bases (*C* and *T*) is approximately equal from base eight to the midpoint of each intron (fig. 2B). From then pyrimidine bases become visibly and steadily more abundant until four bases from the 3’ end of the intron. These last four bases are already part of the splice signal. The length of this pyrimidine-rich tract varies for different lengths of introns, with longer introns having longer pyrimidine-rich tracts. For consistency among length classes, we always extracted a region of 10 bp length from the 3’ end after excluding the last four bases (fig. 3).

By taking the minimal consensus position range among each length class for analyzed regions, we also avoided to include the branch point, which shows significant sequence variability and does not seem in a fixed distance from either the 5’ or the 3’ end.

#### Neutrality of the 5LR

By exploring DNA-strand asymmetry patterns, we aimed to detect whether there are localized, systematic biases for particular motifs in the 3PT. If there are such motifs, the comparison with the presumably neutral 5LR may give insights about the selection strengths for these motifs. However, even in the absence of splicing-coupled selection, neutral processes might create strand asymmetry. And it is not yet clear if the evolution in the 5LR is strand symmetric and different from the 3PT at all. Hence we explored strand symmetry in this region with two tests.

Both tests show that most of the 5LR motifs deviate from the null hypothesis of DNA-strand symmetry (fig. 4A-B). As the null-hypothesis for the chi-squared and equivalence test is identical, trimers with the lowest chi-squared *p*-values unsurprisingly also have the most extreme equivalence test CI.

**Figure 4:**
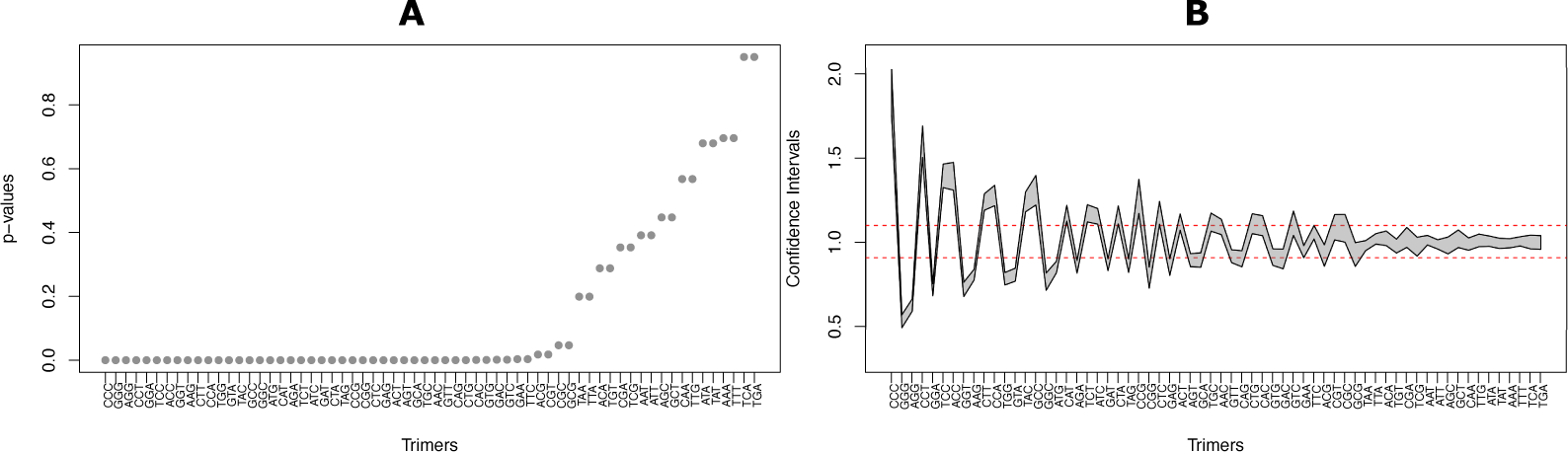
Tests of strand symmetric evolution in the 5LR. (A) Ordered *p*-values from chi-square tests, for the equality of forward and reverse complement trimers. (B) Confidence intervals for the ratios of forward and reverse complement trimers. Red dashed lines correspond to the tolerance range to assume equivalence.

Given the significant deviations from strand symmetry, we can say that the 5LR is not conforming to the strand symmetric evolution. Yet, the more extreme deviations observed for all 3PT motifs (fig. S1A-B), compared to the relatively slight deviations for some 5LR motifs, indicate different causes for asymmetries in these two regions.

Furthermore, despite the utility of the 5LR of short introns as a neutral reference in population genetic analyses, their biallelic frequency spectra deviate from the neutral prediction of symmetry, with an excess of high-frequency *GC* variants, both in autosomes and the X chromosome (fig. S2A-B). Nevertheless, in short introns, evidence for the presence of directional selection is equivocal (Parsch et al., 2010; Halligan & Keightley, 2006; Clemente & Vogl, 2012; Vogl & Bergman, 2015), with no plausible biological explanation for a selective constraint at the base composition level. This directional force favoring *GC* is either explained by context-dependent mutational pattern (Clemente & Vogl, 2012) or by gBGC (Jackson et al., 2017). If gBGC is a meiotic process, we would expect the effects of gBGC to be similar between autosomes and the X chromosome, since the lack of recombination in male meiosis in *Drosophila* and the reduced effective population size of the X chromosome cancel out. This leads to the same expected intensity of gBGC for autosomes and the X chromosome, unlike with directional selection.

Bearing this in mind, we quantified the effect of gBGC for autosomes and the X chromosome from the SFS of *GC*-changing mutations constructed based on their segregating *GC* frequency. We found a weak force, significantly different from *B* = 0, favoring *GC* both in autosomes and X (table 1). On the other hand, the values inferred from the *GC*-conservative mutations are not significantly different from 0, as expected under gBGC. Additionally, to understand whether the strength significantly differs between autosomes and X, we compared the pooled data likelihood to the sum of likelihoods. *B* values inferred separately do not fit the data significantly better than the pooled data (LRT *χ*_*df=1*_^2^ = 2.238*, p* = 0.135). This indicates the same efficiency of the *GC* favoring force in autosomes and the X chromosome, further suggesting gBGC as the cause of the deviation in the SFS, rather than selection.

**Table 1:** *B* values inferred from the SFS of *GC*-changing and conservative mutations, for autosomes, X chromosome and pooled data (autosomes+X). Significance of values are from the LRT, comparing *B* = 0 model.

| | $B$ (*GC*-changing) | $B$ (*GC*-conservative) |
| --- | --- | --- |
| Autosomes | 0.499<sup>***</sup> | -0.041 |
| X | 0.596<sup>***</sup> | -0.026 |
| Pool | 0.51<sup>***</sup> | -0.04 |

<sup>\*\*\*</sup> $p < 0.001$

### Strand asymmetry patterns

Next, we studied strand asymmetry patterns in both regions to examine the selective pressure exerted by the splicing process and observe the differences/similarities between the patterns created by neutral versus selective forces. Initially, we concentrated on the mononucleotide asymmetry. The proportions of complementary nucleotides both in the 5LR and 3PT of autosomal and X-linked introns significantly differ from the 1:1 ratio (table 2, S1). In the 5LR, there is a slight excess of *T* over *A* (*χ*_*df=1*_^2^ = 18.464, *p <* 0.001; *χ*_*df=1*_^2^ = 27.928, *p <* 0.001 for autosomes and the X chromosome, respectively) and a bigger excess of *C* over *G* (*χ*_*df=1*_^2^ = 449.45, *p <* 0.001; *χ*_*df=1*_^2^ = 56.976, *p <* 0.001 for autosomes and X, respectively). The proportion of nucleotides were calculated from the consensus sequences, and are similar to previously reported values in Bergman et al. (2017), calculated from monomorphic sites. The same qualitative pattern is observed for the 3PT, where the bias towards *T* and *C* is much more pronounced (between *A* and *T χ*_*df=1*_^2^ = 14474, *p <* 0.001; χ_*df=1*_^2^ = 1940, *p <* 0.001 for autosomes and X, respectively and between *G* and *C χ*_*df=1*_^2^ = 9877.3, *p <* 0.001; *χ*_*df=1*_^2^ = 1517, *p <* 0.001 for autosomes and X, respectively). Accordingly, the resulting asymmetry scores are higher for the 3PT and *S_CG_* (table 2).

**Table 2:** Proportion of nucleotides (*A, T, G, C*) and mononucleotide asymmetry scores (*S_CG_, S_T A_*) in 5LR and 3PT of autosomal introns

| | $A$ | $T$ | $G$ | $C$ | $S_{CG}$ | $S_{TA}$ |
| --- | --- | --- | --- | --- | --- | --- |
| 5LR | 32.01% | 32.68% | 16.45% | 18.86% | 6.81% | 1.02% |
| 3PT | 21.12% | 46.95% | 8.66% | 23.27% | 45.78% | 37.95% |

The observed nucleotide bias, an excess of the pyrimidines *T* and *C*, conforms to that previously noted for the 3PT, which has been labeled “polypyrimidine” tract. The trend is more subtle but similar in the 5LR. As the 5LR of short introns is considered to be least selectively constrained, the trend there should reflect non-selective processes, *i.e.,* transcription-associated mutation bias. As such processes should be similar over the whole region, we investigated how the trend is changing depending on the position. The 5LR shows similar scores along its length without a significant correlation with position, whereas the biases in the 3PT increase slowly but significantly towards the 3’ junction (fig. 5). It thus seems that the selection pressure is increasing with increasing proximity to the splicing signal.

**Figure 5:**
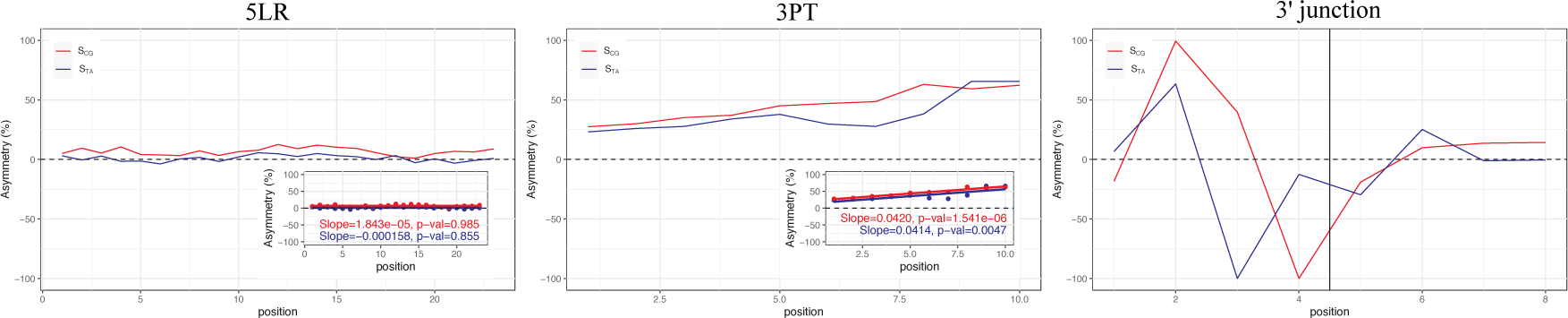
Per position mononucleotide asymmetry scores in 5LR, 3PT and the 3’ junction. *T A* and *CG* asymmetries are shown in blue and red, respectively. Dashed horizontal lines corresponds to symmetry at 0. Vertical line on the far right plot corresponds to the intron-exon boundary. The inset plots for 5LR and 3PT show the regressions between position and asymmetry scores.

If over-representation of pyrimidines in the 3PT is affected by selection against premature splicing, it should not only act on a single base; rather the 3’ splice site consensus sequence is a tetramer, *Y AG|G*, where especially the central *AG* is conserved. Selection should thus suppress the occurrence of this particular motif near the splice junction, such that oligonucleotides associated with the 3’ splice signal should be strongly under-represented on the coding strand in the 3PT compared to the 5LR. The most conserved sequence in the tetrameric splicing signal is the dimer *AG* and it is unclear how much the preceding *Y* and following *G* contribute, *i.e.,* if a dimeric, trimeric, or tetrameric motif is most strongly selected against. We thus explored asymmetries of all possible dimeric, trimeric, or tetrameric motifs. Trimers provide more information compared to dimers, and allow estimation of the relative importance of bases following or preceding the *AG* motif, while the shear number of tetramers makes them difficult to present. We thus show the results for trimers in the text. Generally, results from dimers and tetramers (which can be found in the supplementary information) confirm those from trimers.

Asymmetry scores are symmetrically distributed around 0, since each motif and its reverse complement have the same value with opposite signs. In the 5LR, mononucleotide proportions of complementary bases differ only slightly, such that counts of oligonucleotides and their complement should be approximately equal. As expected, pairs of asymmetry scores from the 5LR are close to symmetry with a relatively low variance of 0.01 for both autosomes and the X chromosome. The variance of pairs of asymmetry scores from the 3PT is much higher with 0.39 for the autosomes and 0.42 for the X chromosome (fig. 6A, S3A). This indicates selection in the 3PT. Tellingly, the most under-represented trimers in the 3PT all contain the *AG* dimer, *i.e.,*— the most conserved part of the consensus 3’ splice site. Indeed, the systematic biases in these motifs, compared to the 5LR and other random motifs in the 3PT, cannot be explained by neutral evolution. Rather they seem to be the result of selection depleting splicing signals from the polypyrimidine tract.

**Figure 6:**
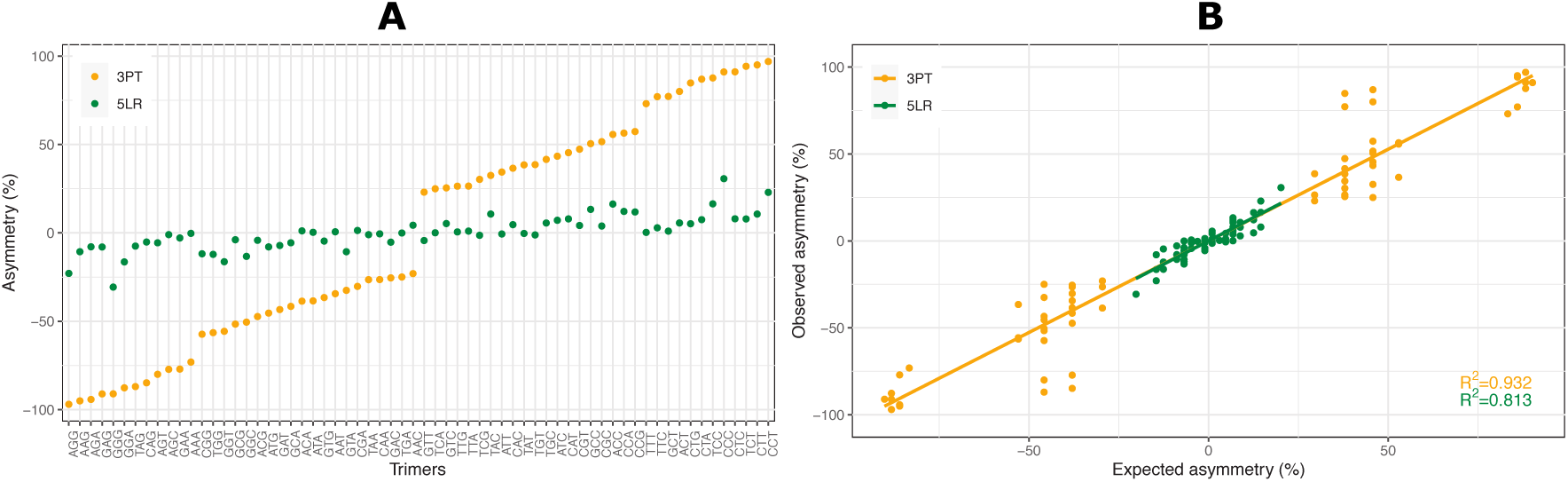
(A) Asymmetry scores per trimer, per region. (B) Correlation between the observed asymmetry of trimers and that expected from the base composition for each region. Orange dots corresponds to 3PT, while green dots are 5LR.

Next, we asked whether there is a correlation between the asymmetries of oligo- and mononucleotides. Because of the limited information of mononucleotides, their asymmetry patterns hardly allow inference of splicing-coupled selection. However, under-representation of particular motifs in the 3PT, together with the position-dependent skew, can be related to avoidance of splicing signal in the 3’ extreme of introns. To understand the connection between the two levels of asymmetries, we examined the relationship between the observed asymmetries of trimers and that expected from the base composition of the region under consideration (fig. 6B). They are highly and significantly correlated (Pearson *R*^2^ = 0.815, *P <* 0.0001; *R*^2^ = 0.932, *P <* 0.0001, for 5LR and 3PT, respectively). In the case of the 3PT, this high correlation suggests that the extreme skew in the base composition (high pyrimidine content) can indeed be affected by selection against the most conserved part *AG* of the 3’ splice signal.

#### Selection coefficients via strand asymmetry

Deviations from strand symmetry in the 5LR can not be associated with selection-driven processes, either at the mono- or oligonucleotide level. The observed slight, constant deviations along the region possibly reflect the effect of transcription-coupled repair leading to mutational asymmetries (transcription-associated mutation bias-TAMB), in line with the recent studies (Bergman et al., 2017; Deger et al., 2019; Törmä et al., 2020). Furthermore, deviations in the biallelic frequency spectra of the 5LR are more compatible with the neutral process *GC*-biased gene conversion rather than directional selection. The inferred strength of directional force causing the deviation is not significantly different from 0 for *GC*-conservative mutations, as opposed to those from *GC*-changing mutations. This supports the presence of a directional force differing between these two classes of mutations, likely gBGC. Additionally, due to the lack of recombination in *Drosophila* males, we expected that the intensity of gBGC should be identical for autosomes and X, and the expectations are met. Thus, even though strand symmetric evolution can not be assumed for the 5LR, it can still be used as a neutral reference for each motif in the 3PT, as the non-adaptive forces affecting these two regions should be similar.

Selection was assessed as the force leading to depletion or excess of motifs in the 3PT compared to the 5LR, which takes into account the presence of non-adaptive forces. As with the asymmetry scores, the strongest negative selection coefficients belong to the trimer motifs containing *AG* (fig. 7, S3B). Strikingly, the three lowest selection coefficients belong to the motifs *AGG*, *TAG*, and *CAG*, perfectly corresponding to the consensus 3’ splice signal, *Y AG|G*. While selection against these trimers is relatively strong with absolute values of nearly three, the highest inferred positive selection coefficient is around 1, much lower than in the reverse direction. These slightly over-represented sequences are rich in pyrimidines, *i.e., C* and *T* . Higher selection coefficients against trimers involved in splicing than those for pyrimidine-rich trimers support further that over-representation of pyrimidines is partly a result of selection against splicing motifs.

**Figure 7:**
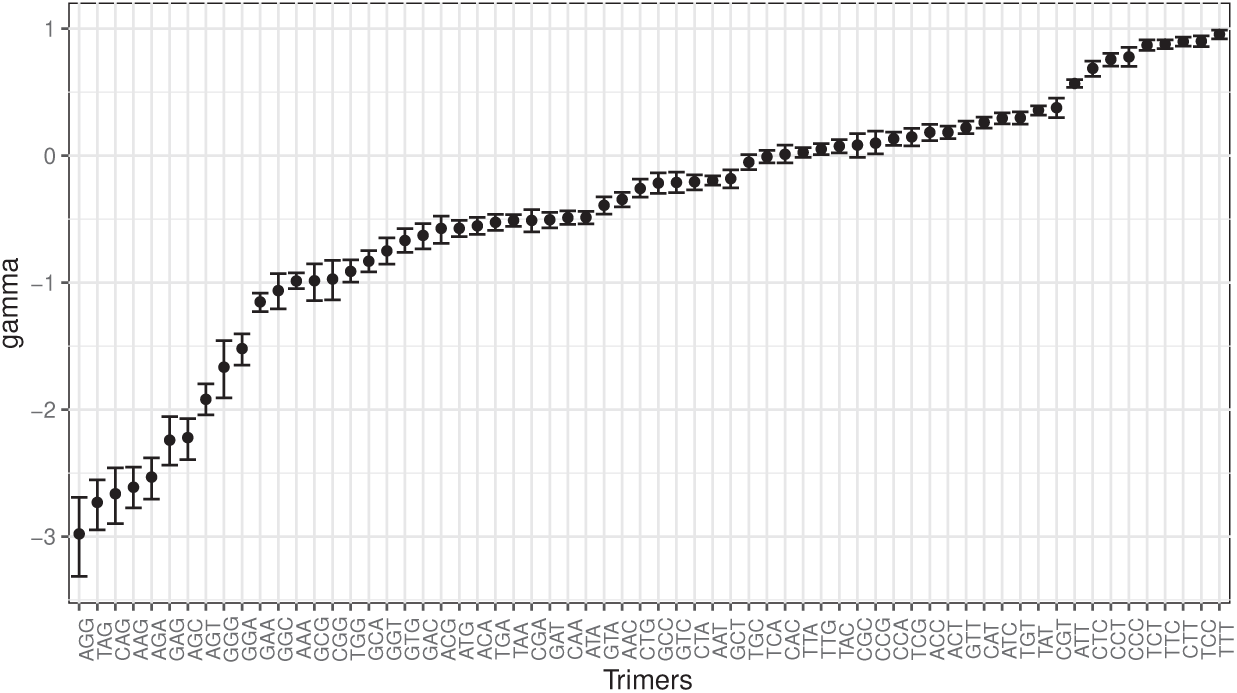
Scaled selection coefficients of each trimer (*γ*(*M*)) in autosomal 3PT, calculated by equation 4. Error bars represent 95% CIs from 1000 bootstraps of the datasets.

The results from the dimer and tetramer analyses confirm this pattern: motifs containing *AG* and *Y AG* have the most extreme skew in the asymmetry patterns and higher selection coefficients (fig. S4). Interestingly, there is a slight increase in the inferred selection strengths from dimer to tetramer. Stronger purifying selection might be expected as the motif gets longer and therefore more specific. Strong selection on long motifs may lead to apparent selection on shorter motifs, even affecting the base composition leading to the over-representation of pyrimidines.

Moreover, we did not observe systematic biases in asymmetry or high selection strengths for the parts of the hexameric 5’ splice signal at the dimer (*e.g., GT*), trimer (*e.g., GTR*, *GGT*) or tetramer (*e.g., GTRA*, *GGTR*) level, but for 3’ splice signal starting from dimers (fig. S4).

As predicted from the sequential nature of splicing, selection against the 3’ splice signal in the pyrimidine tract should be stronger than that against the 5’ splice signal. Indeed, inferred selection strengths against motifs of the 3’ splice signal are always stronger compared to those of the 5’ splice signal (fig. S5). We are aware that the 5’ splice signal also contains the *AG* dimer and has relatively high purine content. However, the most conserved part in the 5’ splice signal is the *GT* dimer, which is not depleted at the dimer level (figs. S5 and S4). Therefore, we posit that strong selection against the 3’ splice signal affects the base composition to create the polypyrimidine tract. Even though we excluded regions overlapping with coding sequences, there are studies showing an excess of phase 0 introns in alternatively spliced genes (Long et al., 1995) and higher sequence conservation of splice signals in exons flanking phase 0 introns (Long & Deutsch, 1999). Thus, to assess whether intron phases influence our estimates, we created additional datasets including (i) only non-phase 0 introns and (ii) only phase 0 introns in *D. melanogaster*. Since common phase 0 introns between *D. melanogaster* and *D. simulans* might also be associated with a higher conservation, we created another dataset (iii) excluding all common phase 0 introns. Selection coefficients were similar for each trimer in all three datasets (fig. S6), showing that the magnitude of selection varies little within the genome.

We also analyzed the introns of *D. simulans* (fig. S7). The magnitude of the selection strength is similar and consistent for each trimer motif in both species. Although the splicing machinery is highly conserved and we would expect selection against the splicing signal in the polypyrimidine tract to be similar among species, it should be noted that the similar patterns between these two *Drosophila* species may not necessarily be due to a universal mechanism in intron evolution. In fact, the majority of sites are shared between the two species being compared, suggesting that the observed patterns may be the result of phylogenetic inertia. To explore evolutionary universality, we rather utilized the asymmetry patterns in other eukaryotic species in the last section.

#### Positional effect on the base composition of the 3PT

In the light of previous studies, we proposed three hypotheses that might explain the base composition evolution in 3PT: hypothesis one (HI) posits selection is only for pyrimidines, hypothesis two (HII) posits only avoidance of *AG* dimer, and hypothesis three (HIII) posits both selection for pyrimidines and selection against *AG*. In the previous section, we systematically compared asymmetry patterns of all dimers, trimers, and tetramers in the 3PT to those in the 5LR. Results indicate that 3’ splice signal associated motifs are most strongly underrepresented, which supports HII. One might argue that selection for pyrimidines (HI) can create under-representation of motifs rich in purine bases, however this cannot explain the stronger avoidance of the trimer *TAG* compared to *GGG* (fig.7), since the former contains two purines and the latter three. Rather this gives HII an advantage over HI, yet HIII may still be preferable to both HI and HII.

For comparing the three hypotheses quantitatively, we applied a method to infer the selection strength under these three models and compared their likelihoods. Selection was modeled through its effect on fixation probabilities (eq. 5) and given the mutation matrix at neutral equilibrium inferred from the 5LR and the joint frequency matrix of four bases at position *i* and *i*+1 (1 *≤ i ≤* 9) in the 3PT, we obtained the selection coefficients for each position. Under the first hypothesis, we obtained selection strengths for all monomers (*A, T, G, C*). For the second hypothesis only *AG*, and for the third hypothesis both monomer and *AG* coefficients were inferred. We incorporated the selection against *AG* only through the positional effect, meaning that selection is conditional on an *A* preceding or a *G* following the focal position. Normally, selection on dimers and on higher-order oligonucleotide motifs associated with the 3’ splicing signal would create a selective effect on the monomer level. Yet it is hard to disentangle the effect of dimer selection on the monomer level or to consider selection on every possible oligonucleotide. Thus, by inferring only the *AG* selection coefficient for the HII, we only test the effect of sequence context in the simplest scenario.

Comparison between the models with likelihood ratio tests reveals that both for autosomes and the X chromosome, HIII fits significantly better than both HI and HII at all positions of the 3PT (tables S2 and S3). This shows that only selection on monomers is not enough, but the sequence context, more specifically selection against the *AG* dimer is necessary to explain the observed patterns. The selection coefficients inferred under the best fit model HIII show that the constraint on the *AG* dimer is stronger than on monomers (figs. 8, S8). However, as we mentioned earlier, the selection on dimer and monomer level is not independent. This is because the longer the sequence, the more information there is by a factor of four per added basepair. Therefore, the selection strength is hard to compare between monomers and dimers. The relative selection strength between pyrimidine bases, *C* and *T*, is roughly similar and varies with the position along the 3PT. In general, selection coefficients of *AG* and monomers increase towards the 3’ end.

**Figure 8:**
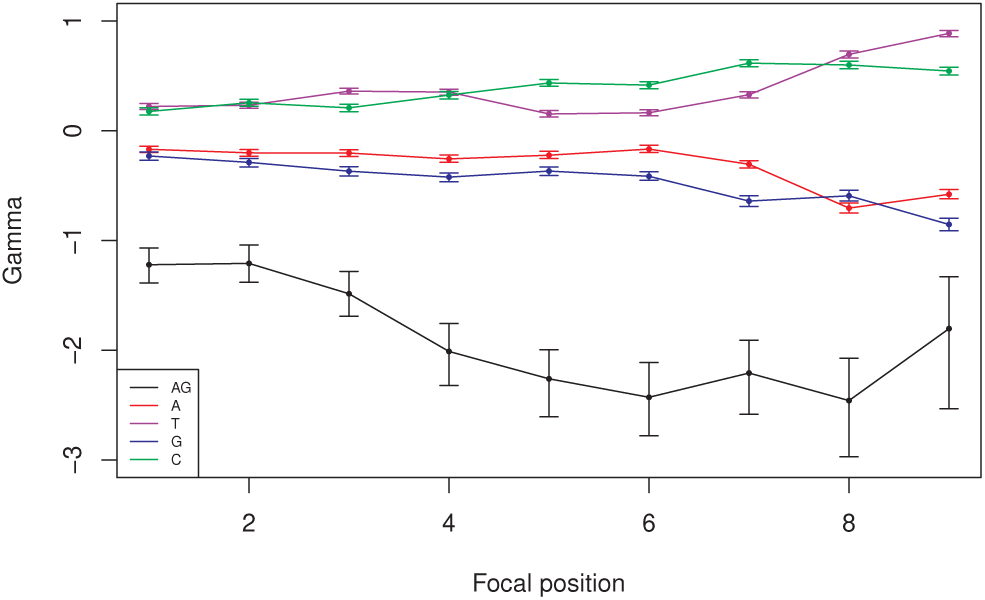
Inferred scaled selection coefficients *γ* of the monomers and dimer *AG* for each position in the autosomal 3PT under the best fit model HIII. Error bars represent 95% CIs from 1000 bootstraps of the datasets.

We validated our inference by simulating a DNA sequence of the same length as the 3PT under the three models. To avoid differences between positions, the sequence was circularly joined such that the first and last positions were adjacent to each other (see Methods). We calculated the joint frequency matrix of the four bases at neighboring sites and used a *χ*_*df=1*_^2^ -inspired statistic to visualize the deviation of the simulated values from the empirical data. Once again the HIII model provides the best fit to the data, as evidenced by the overall lowest values obtained. Moreover, this analysis allowed us to interpret the patterns that each model failed to explain, which might hint at more complex scenarios that were not considered. The HI model, which considers only selection on monomers, fails to explain not only the under-representation of the *AG* dimer but also other motifs, especially in the beginning of the polypyrimidine tract (figs. 9, S9). While selection on both monomers and *AG* (HIII) provides a better fit for motifs that HI cannot account for, it falls short explaining the high frequency of motifs rich in pyrimidines, such as *TT* . Lastly, the model considering only selection on the *AG* dimer shows the worst fit for motifs high in pyrimidine in the beginning of the sequence, and it also deviates for the *AG* dimer towards the end.

**Figure 9:**
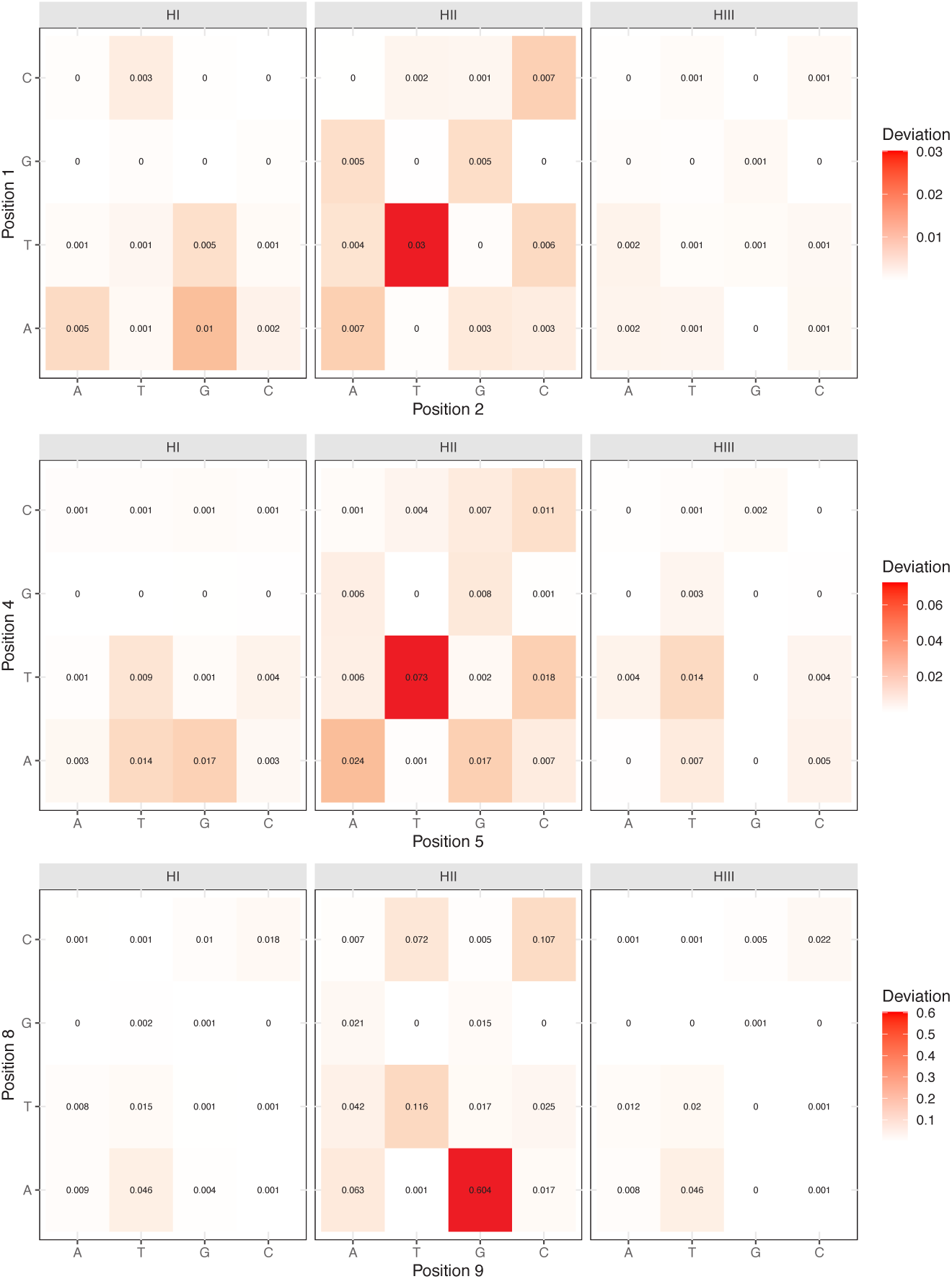
Deviations of the three hypotheses from the autosomal empirical joint frequency data of the four bases. Values calculated as *χ*_*df=1*_^2^ statistic and high to low deviation is represented with red to white colour gradient. Matrices for three positions are chosen to visualize the pattern along the 3PT .

#### Asymmetry patterns of other eukaryotic species

We used the 5LR of *Drosophila* short introns as a neutral reference to compare the DNA-asymmetry patterns in the 3PT, validate the effect of selection, and quantify its strength. While we are not aware of a similar neutrally evolving region in other eukaryotic species, we nevertheless qualitatively checked for the universality of avoidance of the splice signal in the 3PT in other eukaryotes. If 3’ splice signal-associated motifs are similarly under-represented in other eukaryotes, the same evolutionary process is likely at work as in *Drosophila*.

To test this, we analyzed the asymmetry patterns in polypyrimidine tracts of human, sea urchin, worm (*Caenorhabditis elegans*), rice, mouse-ear cress (*Arabidopsis thaliana*), moss, and the two yeast species *Saccharomyces cerevisiae* and *Lachancea thermotolerans*. Comparative analyses suggest that the strength of the polypyrimidine tract with respect to length and pyrimidine content differs between plants, metazoa, and fungi; it is very weak in fungi and shows a gradual increase from plants to metazoa and from *C. elegans* to human within metazoa (Schwartz et al., 2008). The species we selected cover this range of diversity. Among them, *C. elegans* displays unusual properties regarding splicing (Riddle et al., 1997): introns are exceptionally short, most under 60 nucleotides, and seem to lack an obvious polypyrimidine tract. Additionally, even though *AG* is generally highly conserved, they use splicing signals not containing *AG* more often than other species (Riddle et al., 1997). It has been demonstrated that *AG* is not obligatory for the 3’ splice site recognition, since mutations from *G* to *A* or to *C* did not affect splicing (Aroian et al., 1993; Zhang & Blumenthal, 1996). The fungal genomes have also relatively short and few introns (in the hundreds), while other eukaryotes contain thousands of introns (Neuveglise et al., 2011). Moreover, the protein U2AF1, which is a nearly universal part of the spliceosome, is lost in *S. cerevisiae*, but not in *L. thermotolerans* (Neuveglise et al., 2011; Hooks et al., 2014). U2AF1 recognizes and binds to the 3’-AG motif after scanning the region downstream of the branch point, *i.e.,* the polypyrimidine tract (Smith et al., 1993). Comparing these two yeast species gives us a chance to test whether trans-splicing factors associated with this region have a significant effect on the base composition and motif representation in the polypyrimidine tract.

We again focused on the short intron class of each organism to minimize selection not related to splicing, except for the yeast species where all introns were utilized, as they have very few and relatively short introns (fig S10). Even though the relative nucleotide composition varies between species, pyrimidines are over-represented close to the 3’ ends of short introns in all species. In human and sea urchin both *C* and *T* increase in frequency, in the other species only *T* (fig. S11). The distribution of asymmetry scores of dimeric and trimeric motifs in these pyrimidine-enriched regions resembles that in *Drosophila* (figs. 10, S12, S13). At the dimer level, the most under-represented motif is *AG* for all species, except for *C. elegans*. In *C. elegans AA* is the first and *AC* the third most under-represented dimers. Mutational analyses showed these motifs can serve as the 3’ splice site in *C. elegans* (Aroian et al., 1993; Zhang & Blumenthal, 1996). Thus observing them as the most under-represented dimers in asymmetry scores still supports the effect of avoidance of premature splicing in the nucleotide composition evolution of the 3’ region of short introns. At the trimer level, *AG*-containing motifs have again the lowest asymmetry scores in human, sea urchin, rice, and *Arabidopsis*. In moss, *C. elegans* and yeasts, they are also under-represented (negative asymmetry scores), yet do not always have the lowest values. This is in line with the previous studies reporting relatively weaker polypyrimidine tracts and possibly reflects the variation in splicing signal usage in these species (Schwartz et al., 2008).

Recently, Schirman et al. (2021) reported that the absence of the U2AF1 protein in certain yeast species, including *S. cerevisiae*, leads to increased selective pressure to avoid a premature splice signal. In our study, the pattern of asymmetry scores looks grossly similar in *S. cerevisiae* and *L. thermotolerans*. At the dimer level, both species exhibit a strong under-representation of the *AG* motif (fig. 10, S12), and at the trimer level, most motifs containing *AG* have similar asymmetry scores, except for *CAG*, which does not have a high asymmetry score in *L. thermotolerans* (fig. 10, S13). The reason for this seems to be an under-representation of the reverse complement of the *CAG* motif (*i.e., CTG*). We cannot speculate why *CTG* is depleted. A difference between our study and that of Schirman et al. (2021) may be the strategy used to identify depletion of 3’ splice signal motifs. They did not use strand asymmetry but compared the sequence context around introns’ 3’ ends to that of 1000 randomly selected genomic loci and did not investigate depletion of the motifs *TAG* and *CAG* separately, but together. In any case, a systematic depletion of the *AG* dimer and *AG*-containing trimers in both species suggests that comparable base composition and asymmetry patterns can arise even in the absence of the trans-splicing factor U2AF1.

**Figure 10:**
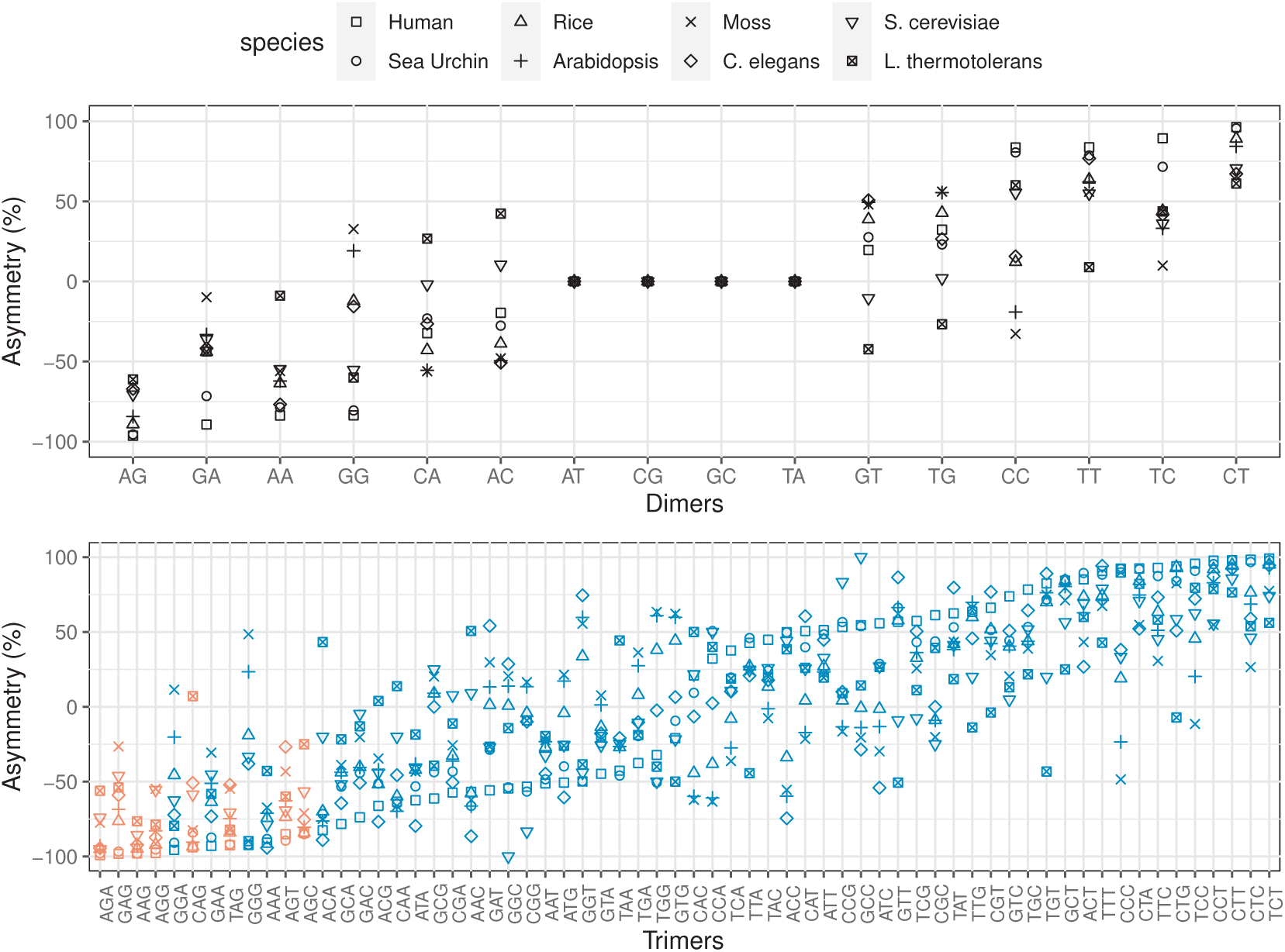
Asymmetry scores of dimers (top row) and trimers (bottom row) from the pyrimidine enriched regions in the introns of 8 eukaryotic species. Species are shown with different shapes and trimer motifs containing *AG* in it are depicted with red symbols, while non-*AG* motifs are represented with blue.

## Discussion

Many *in vivo* and *in vitro* experiments have shown the functional importance of the 3PT in splicing (Green, 1986; Spellman et al., 2005). Indeed, the 3PT serves as the binding site for trans-splicing factors (Singh et al., 1995) and mutations in it might result in a decreased splicing efficiency. Although it is well-known that the nucleotide composition of the 3PT varies among introns and species, its co-evolution with the trans-splicing factors is still largely unknown. The belief that the composition of the 3PT evolved due to selective preference of pyrimidines has been challenged by studies showing that the presence of purines is not detrimental to splicing (Roscigno et al., 1993) and that splicing factors binding to the polypyrimidine tract can tolerate different base compositions (Singh et al., 1995, 2000). Thus selection at the monomer level generally favoring pyrimidines, especially uridines, over purines may be rather weak. On the other hand, strong selection against premature splicing may lead to avoidance of the strongly conserved splice signal *AG*, during scanning mechanism from the branch point towards the 3’ end of the intron, to avoid premature splicing. This selection may also contribute to the base composition evolution on the monomer level. Generally, our results show that the high pyrimidine content is the result of purifying selection against spurious or cryptic 3’ splice signals, thus against *AG*, as well as the selection for pyrimidine bases, likely due to the binding affinities of trans-splicing factors.

Fruit flies of the genus *Drosophila* have mainly short introns. Within these short introns, the function of the 5LR between the 5’ splice signal and the branch point seems to be mainly or exclusively to form a loop of the required length. Consequently, sequence evolution within the 5LR appears to be unconstrained and has been used as neutral reference for inference of selection (Parsch et al., 2010; Lawrie et al., 2013; Machado et al., 2020). We also found no association of the nucleotide composition in the 5LR with selective processes both in population genetic analyses of site frequency spectra and DNA-asymmetry patterns. On the other hand, the sequence of the 3PT is functionally important for splicing. Therefore, we used strand asymmetry patterns to identify and quantify selection in the 3PT by comparing it to 5LR. We find that conserved motifs in the 3’ splice site are more avoided than others with similar or higher pyrimidine content, from which we conclude that selection in the 3PT is not exclusively for pyrimidines.

To infer the relative importance of avoidance of the canonical splice site and selection for higher pyrimidine content, we compared the fit of three models to the data using a newly developed inference method accounting for positional effects along the 3PT: I) selection for pyrimidines, II) selection against the *AG* dimer or III) selection both for pyrimidines and against the *AG* dimer. Our results show that both selection for pyrimidines and avoidance of the canonical 3’ splice signal are necessary to explain the base composition of the 3PT. Although our method has some limitations, such as not accounting for selection against higher-order oligonucleotides or correlation between selection at dimer and monomer levels, inferred joint frequencies closely fit the empirical data.

In *Drosophila*, the presence of an established neutral reference, the 5LR, enabled the detection of presumably selected motifs by comparing asymmetry patterns and the quantification of selection strength in the 3PT. In other eukaryotes, the preferred splice signal and the nucleotide composition varies, as does the length of the 3PT (Nguyen & Xie, 2019; Coolidge et al., 1997), although pyrimidines are generally over-represented (Coolidge et al., 1997). Due to the lack of a neutral reference in other eukaryotes, a similar approach to quantification of the selection strength is not possible. Nevertheless the asymmetry pattern within the 3PT also suggests the same mechanism as in *Drosophila*: oligonucleotides containing the 3’ splice signal are under-represented in the 3PT of short introns, *i.e.,* the region between the branch point and the 3’ splice signal. Thus selection against premature splicing in the 3PT seems to be universal among eukaryotes.

In a recent study, Rong et al. (2020) proposed an exaptation mechanism (Gould & Vrba, 1982) to explain the evolution of exonic splicing enhancers (ESEs): precursor ESEs would be created by the joint action of mutation bias and purifying selection on the protein code. Once ESE motifs started to appear, ESEs and trans-splicing factors would co-evolve due to selection on splicing. A study in yeast suggested that the avoidance of cryptic splicing through depletion of the 3’ splice signal around the 3’ end drives intron sequence evolution and splicing factors can co-evolve with this (Schirman et al., 2021). In the light of our results, a positive feedback loop is conceivable: once the pyrimidine content in the 3PT increased due to avoidance of the 3’ splice signal, splicing factors co-evolved to develop higher binding specificity to pyrimidines. In any case, the co-evolution of splicing avoidance and binding specificity of trans-splicing factors seems to determine the architecture of short introns, where the splicing information is mainly intronic (“intron-definition”) (Talerico & Berget, 1994).

## Acknowledgements

The authors thank all members of the Vienna Graduate School of Population Genetics for support and discussion. We also thank two anonymous reviewers for their helpful comments on the manuscript. This work was supported by the Austrian Science Fund (FWF; W1225-B20).

---

**Figure S1:**
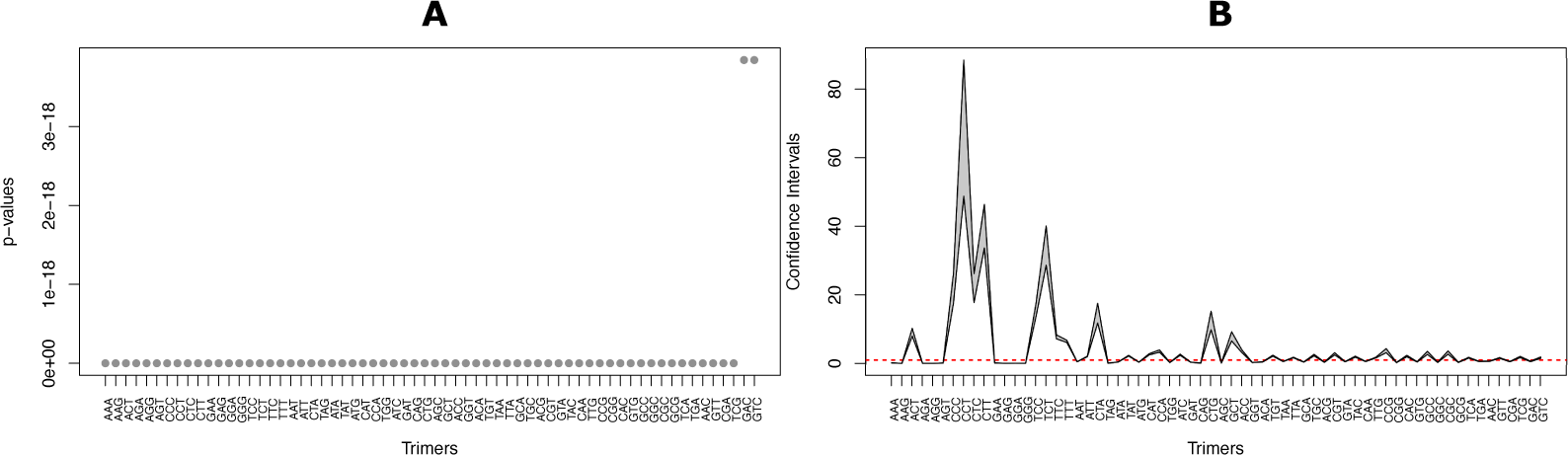
Tests of strand symmetric evolution in the 3PT. (A) Ordered p-values from chi-square tests, for the equality of forward and reverse complement trimers. (B) Confidence intervals for the ratios of forward and reverse complement trimers. Red dashed lines correspond to the tolerance range to assume equivalence.

**Figure S2:**
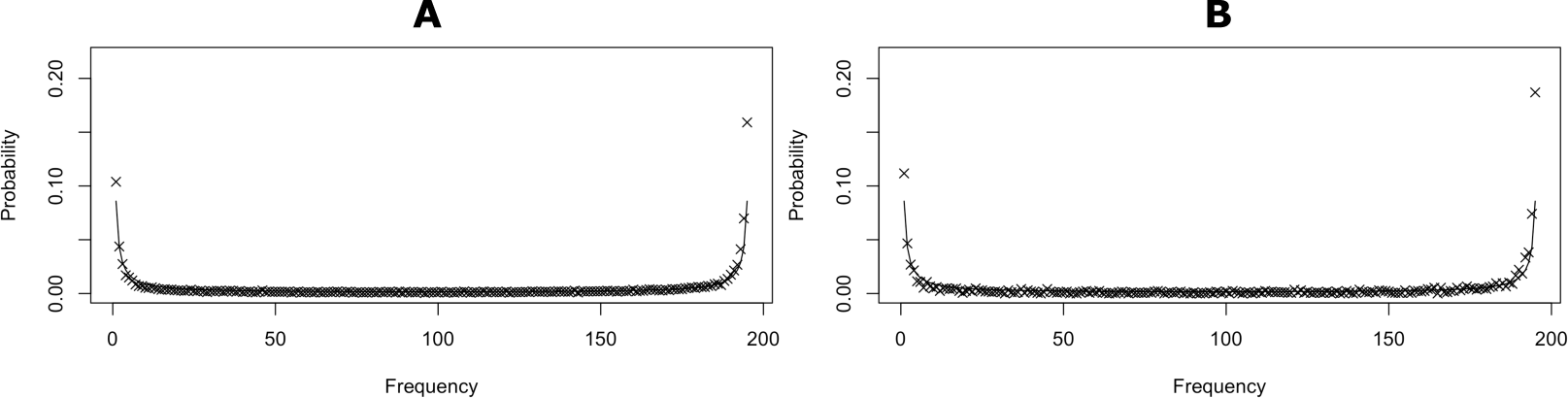
Biallelic spectra of GC-changing mutations (*A/C*, *A/G*, *T/C*, *T/G*) constructed based on their segregating *GC* frequency for (A) Autosomal short introns (B) X-linked short introns. For this analysis, only introns shorter than 66 bp were used. The lines represent expected site frequencies under neutral equilibrium; crosses represent observed site frequencies. The deviations are significant (*χ*_*df=1*_^2^ = 1875.3, *p <* 0.001 and *χ*_*df=1*_^2^ = 518.81, *p <* 0.001 for Autosome and X, respectively)

**Figure S3:**
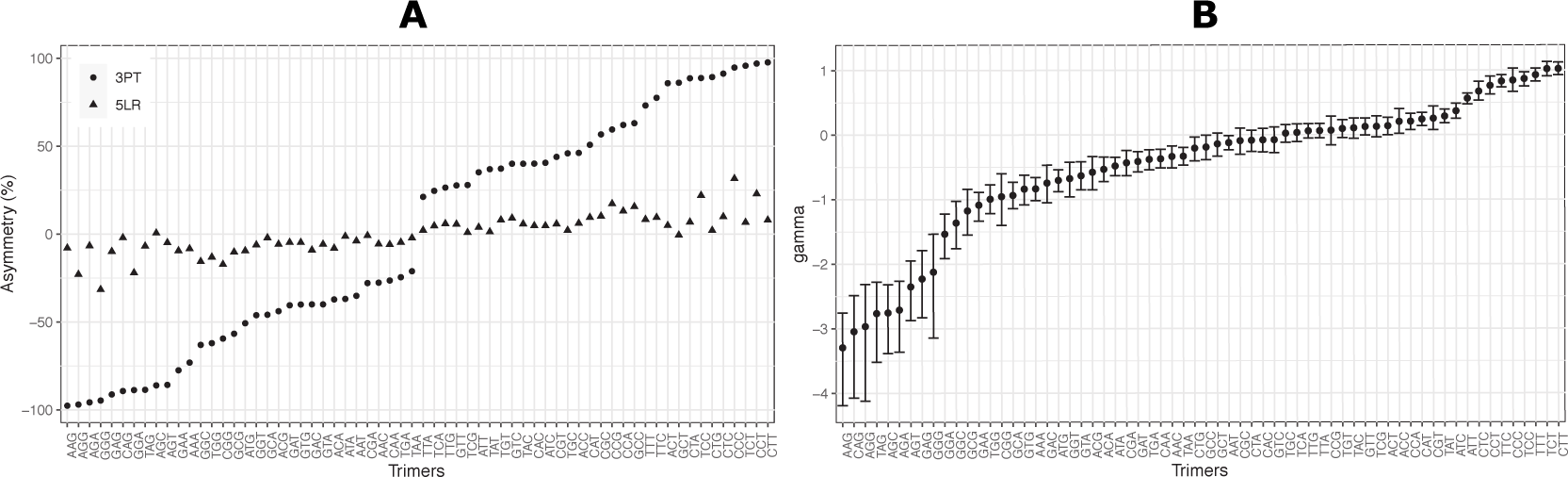
Results for X-linked introns. (A) Asymmetry scores per trimer, per region. Circles represent 3PT and triangles represent 5LR (B) Selection coefficients of each trimer in 3PT. Error bars represent the 95% CIs from 1000 bootstraps of the datasets.

**Figure S4:**
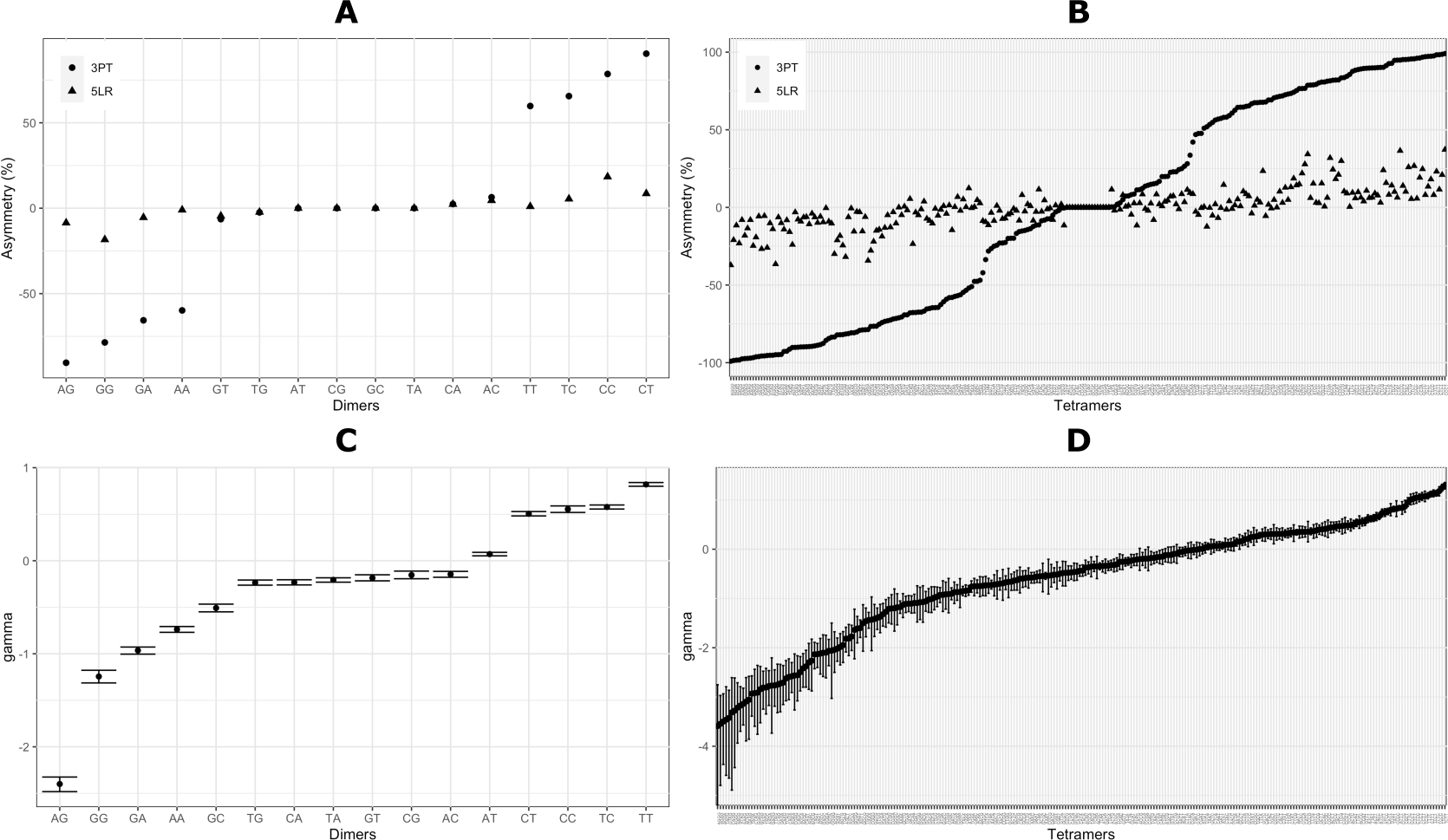
Results for dimers (A-C) and tetramers (B-D). (A-B) Asymmetry scores per region, per dimer and per tetramer, respectively. Circles represent 3PT and triangles represent 5LR (C-D) Scaled selection coefficients of each dimer and tetramer in the 3PT, respectively. Error bars represent the 95% CIs from 1000 bootstraps of the original datasets.

**Figure S5:**
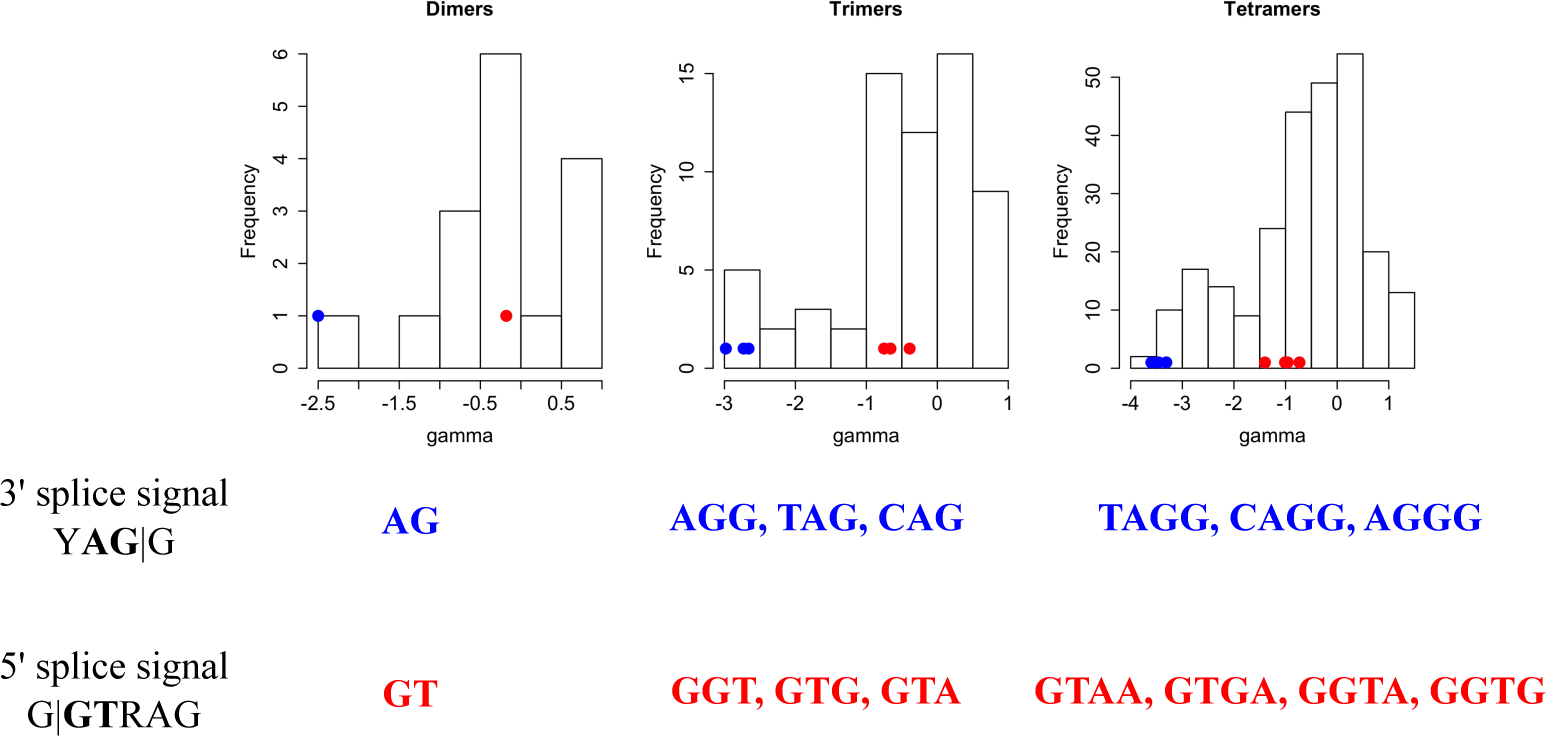
Scaled selection strength (*γ*(*M*)) distributions for dimer (16), trimer (64) and tetramer (256) motifs. Dots in the graphs correspond to the gamma values for specific motifs from 3’ splice signal (blue) or 5’ splice signal (red). The lower panel shows the pinned motifs in histograms.

**Figure S6:**
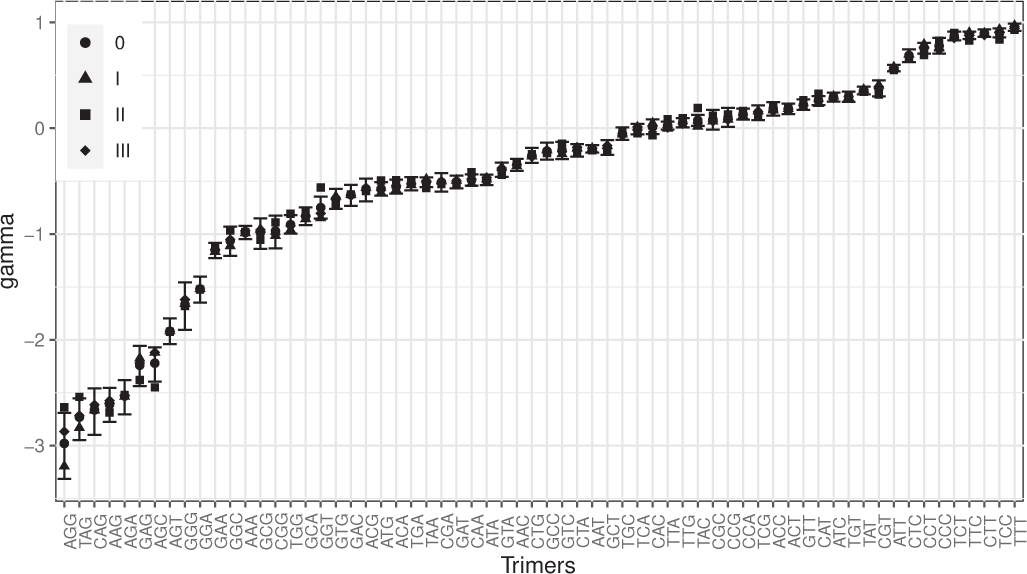
Scaled selection coefficients of each trimer (*γ*(*M*)) in the 3PT for different datasets. 0 (circle): original dataset with all short introns. I (triangle): dataset including only non-phase 0 (3n+1, 3n+2) introns. II (square): dataset including only phase 0 (3n) introns. III (diamonds): dataset excluding the common phase 0 introns between *D. melanogaster* and *D. simulans*. Error bars represent the 95% CIs from 1000 bootstraps of the original datasets.

**Figure S7:**
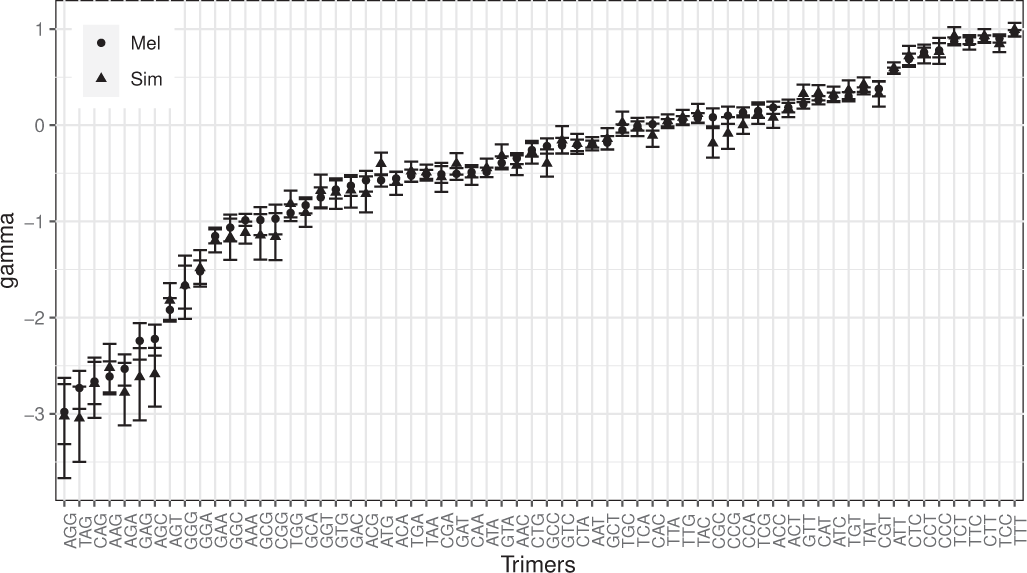
Scaled selection coefficients of each trimer (*γ*(*M*)) in the 3PT for different species. Circles are estimates from *D. melanogaster*, while triangles are estimates from *D. simuans*. Error bars represent the 95% CIs from 1000 bootstraps of the datasets.

**Figure S8:**
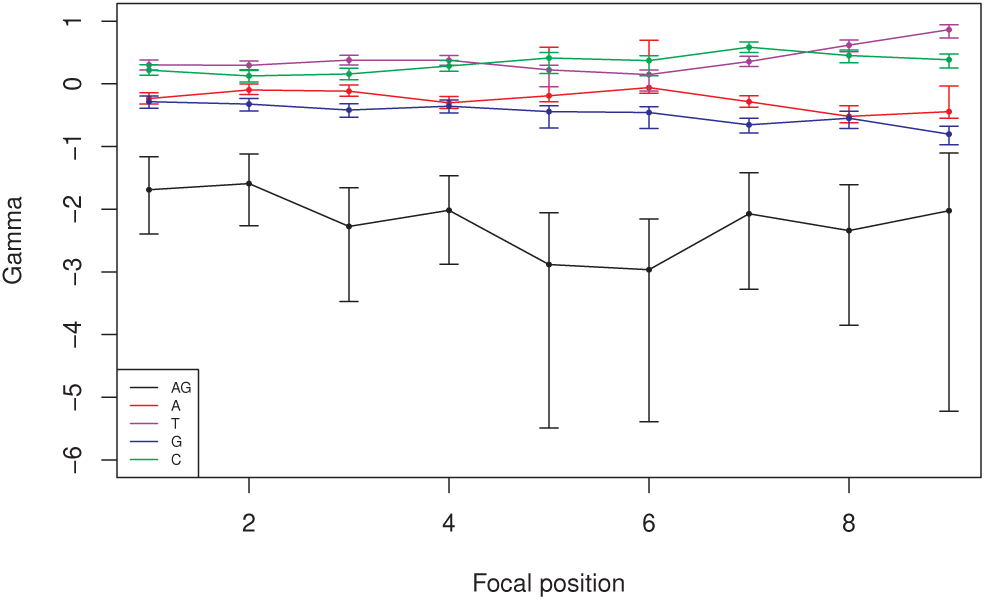
Inferred scaled selection coefficients *γ* of the monomers and dimer *AG* for each position in the X-linked 3PT under the model HIII. Error bars represent 95% CIs from 1000 bootstraps of the datasets.

**Figure S9:**
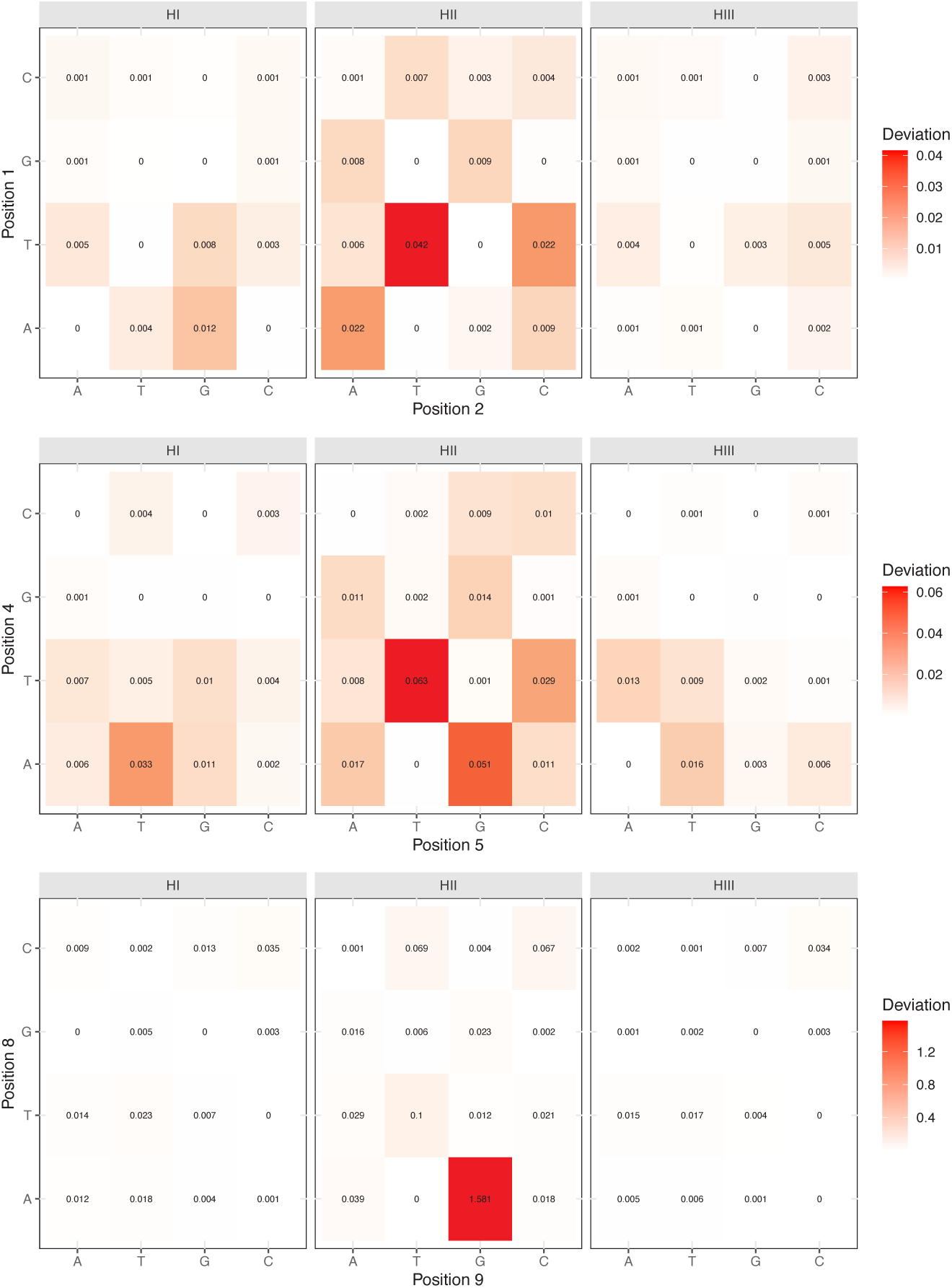
Deviations of the three hypothesis from the X chromosomal empirical joint frequency data of the four bases. Values calculated as *χ*_*df=1*_^2^ statistic and high to low deviation is represented with red to white colour gradient. Matrices for three positions are chosen to visualize the pattern along the 3PT .

**Figure S10:**
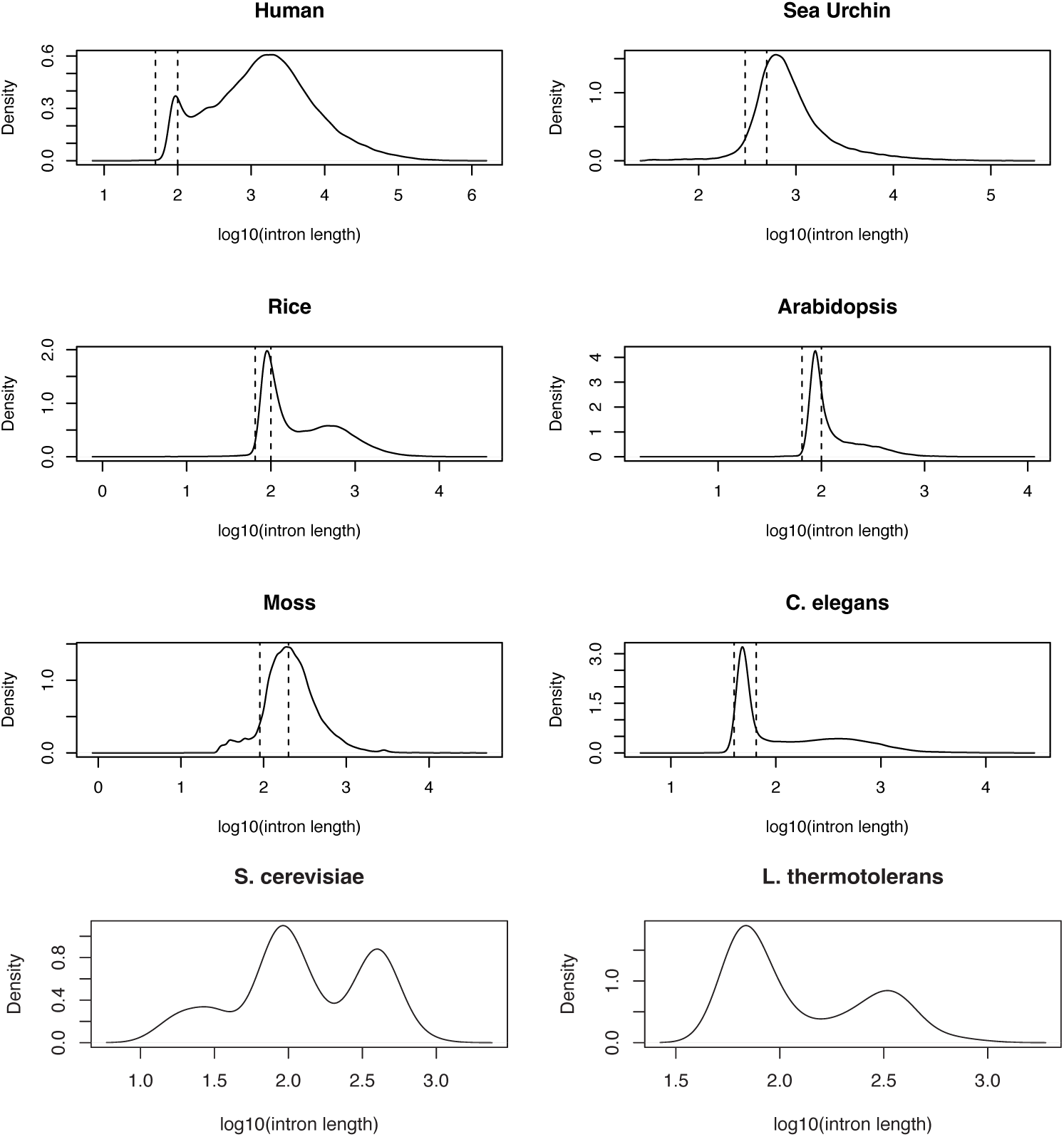
The length distribution of introns obtained for eight eukaryotic species. Vertical dashed lines represent the length range of extracted and analyzed short intron class. Human: 50-100 bp, Sea Urchin: 300-500 bp, Rice: 65-100 bp, *Arabidopsis*: 65-100bp, Moss: 90-200 bp, *C. elegans*: 40-65 bp. For yeast species all annotated introns were utilized.

**Figure S11:**
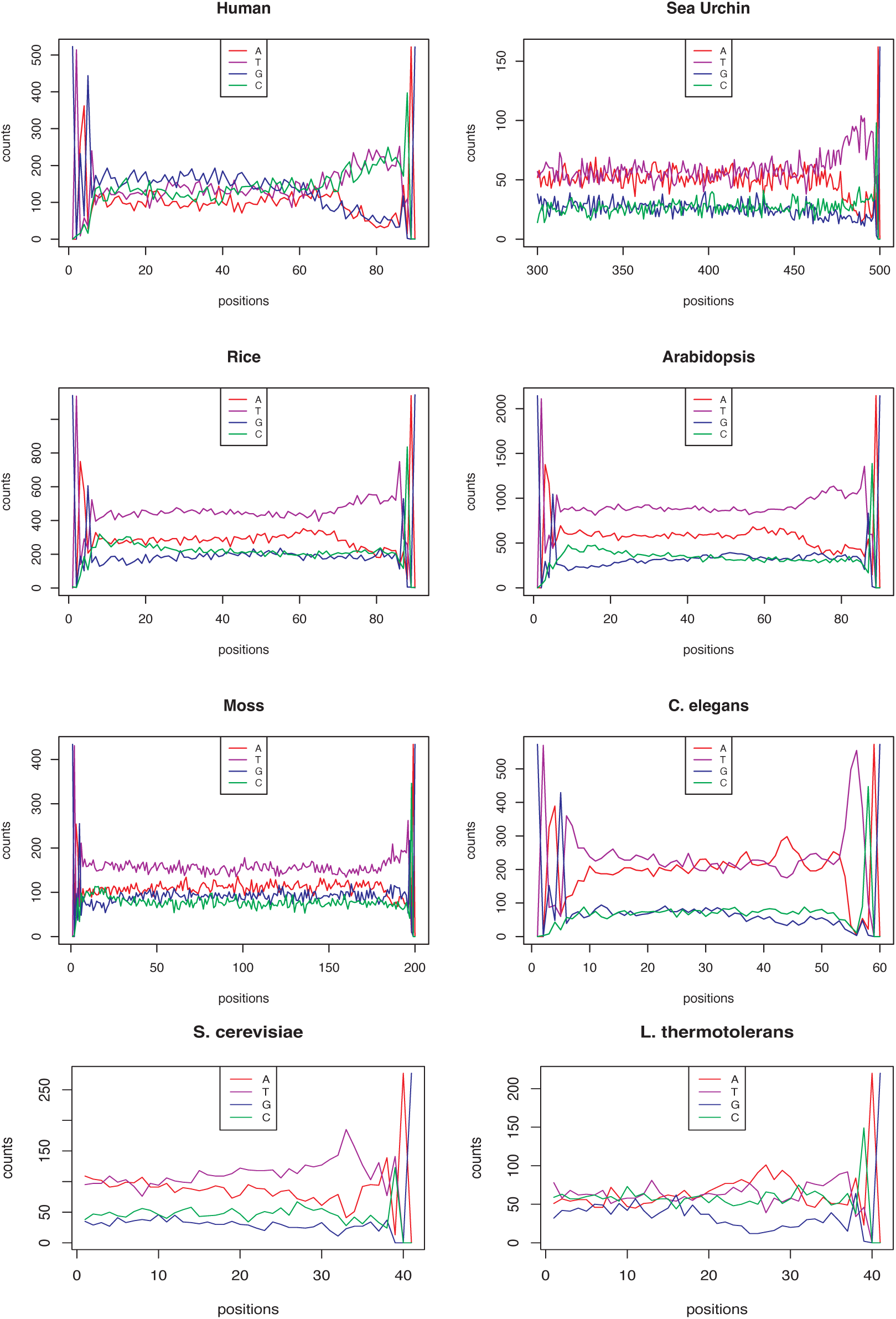
Nucleotide composition of introns with length 90 for human, 500 for sea urchin, 90 for rice and *Arabidopsis*, 200 for moss, and 60 for *C. elegans*. Only one candidate length class were chosen to visualize pattern. Total numbers of introns used for each species are 523, 162, 1152, 2145, 434 and 573, respectively. For yeast species all annotated introns with different lengths were utilized (277 and 220 introns for *S. cerevisiae* and *L. thermotolerans*, respectively), thus nucleotide composition is given for the last 40 bases from the 3’ end.

**Figure S12:**
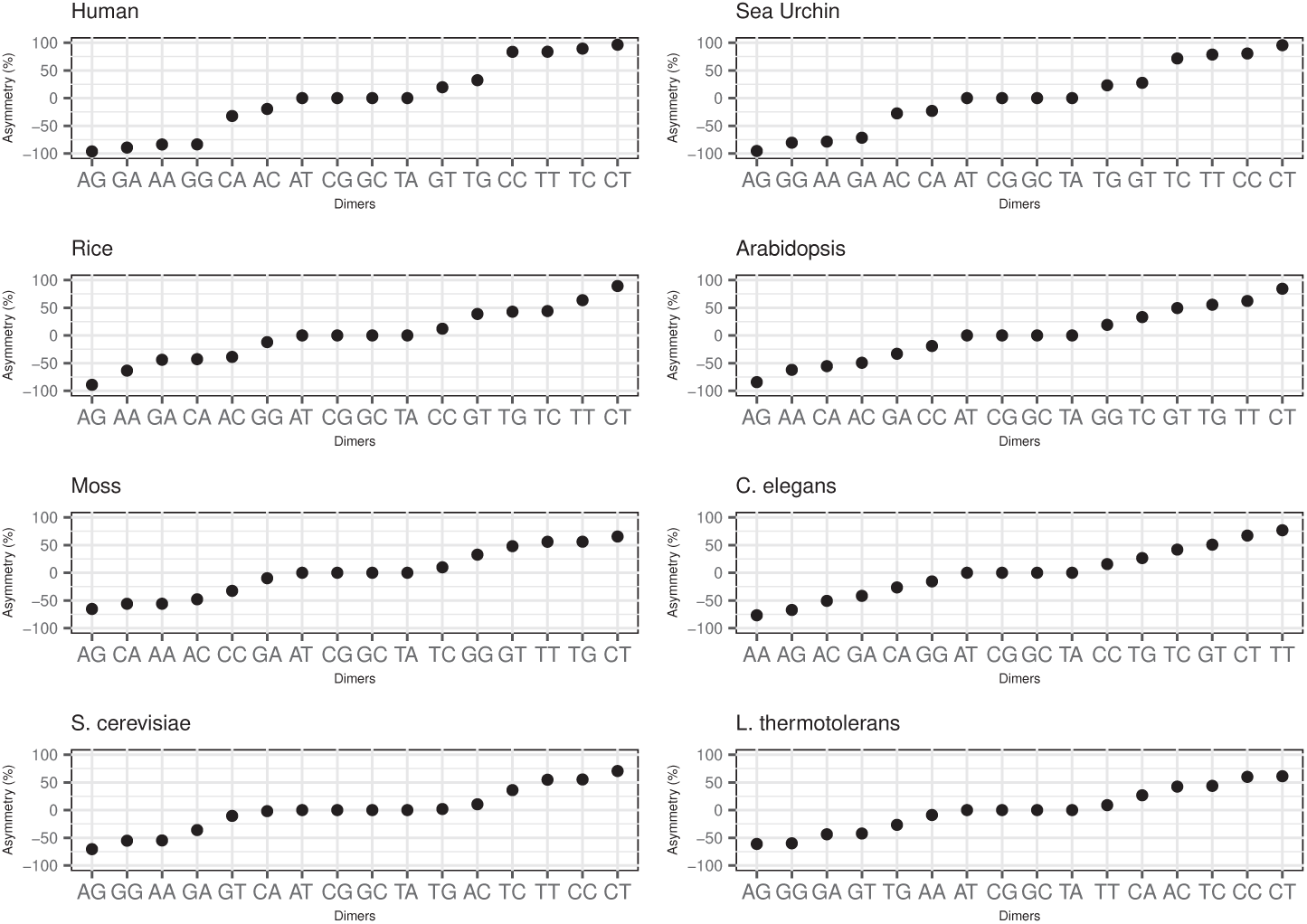
Asymmetry scores of dimers from the pyrimidine enriched regions in the introns of 8 eukaryotic species.

**Figure S13:**
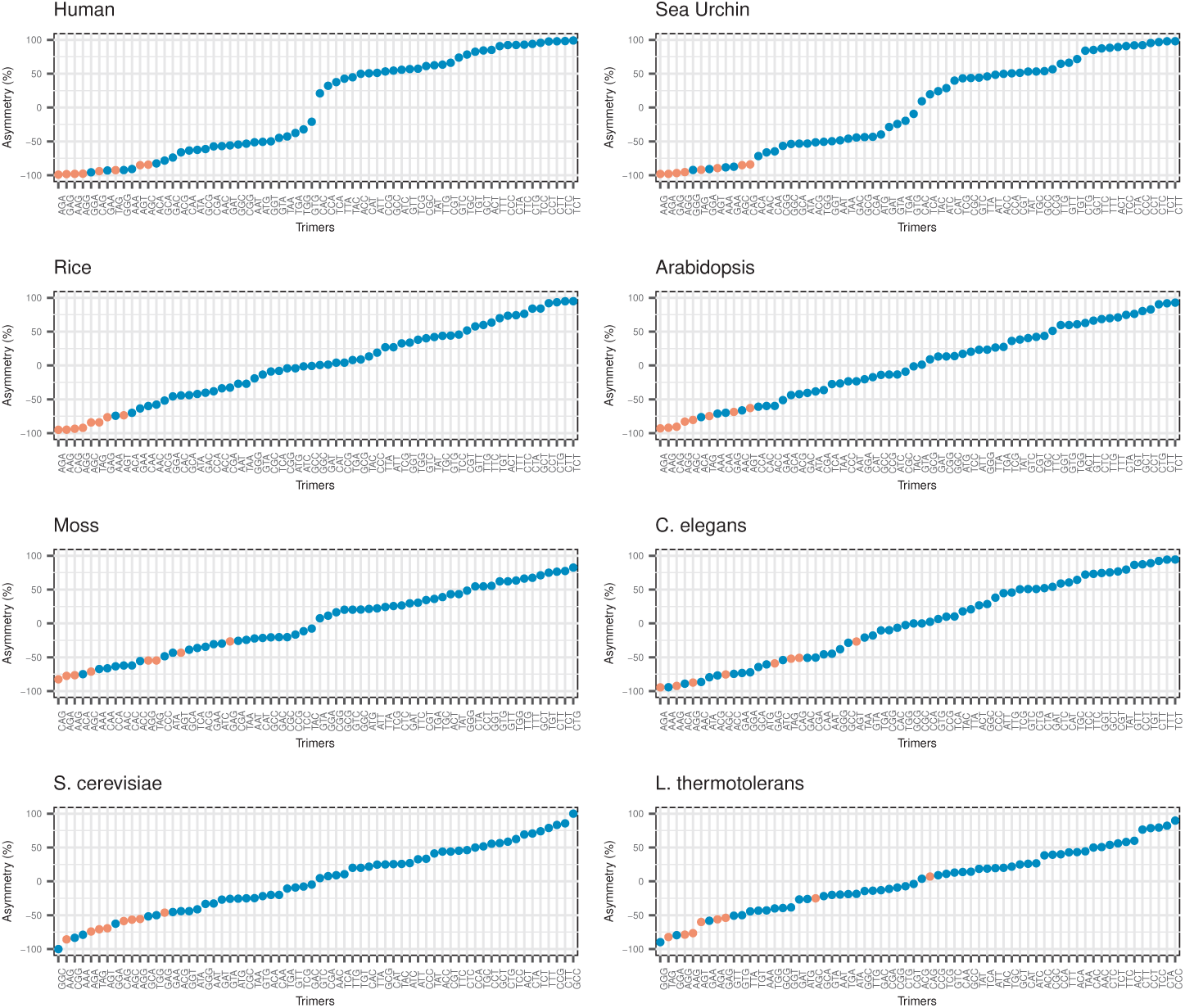
Asymmetry scores of trimers from the pyrimidine enriched regions in the short introns of 8 eukaryotic species. Motifs containing *AG* in it are depicted with red points, while non-*AG* motifs are represented with blue.

**Table S1:**
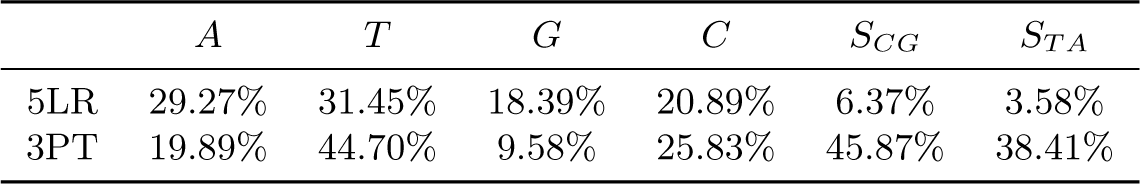
Proportion of nucleotides (*A, T, G, C*) and mononucleotide asymmetry scores (*S_CG_, S_T A_*) in 5LR and 3PT of X-linked introns

| | *A* | *T* | *G* | *C* | $S_{CG}$ | $S_{TA}$ |
| --- | --- | --- | --- | --- | --- | --- |
| 5LR | 29.27% | 31.45% | 18.39% | 20.89% | 6.37% | 3.58% |
| 3PT | 19.89% | 44.70% | 9.58% | 25.83% | 45.87% | 38.41% |

**Table S2:**
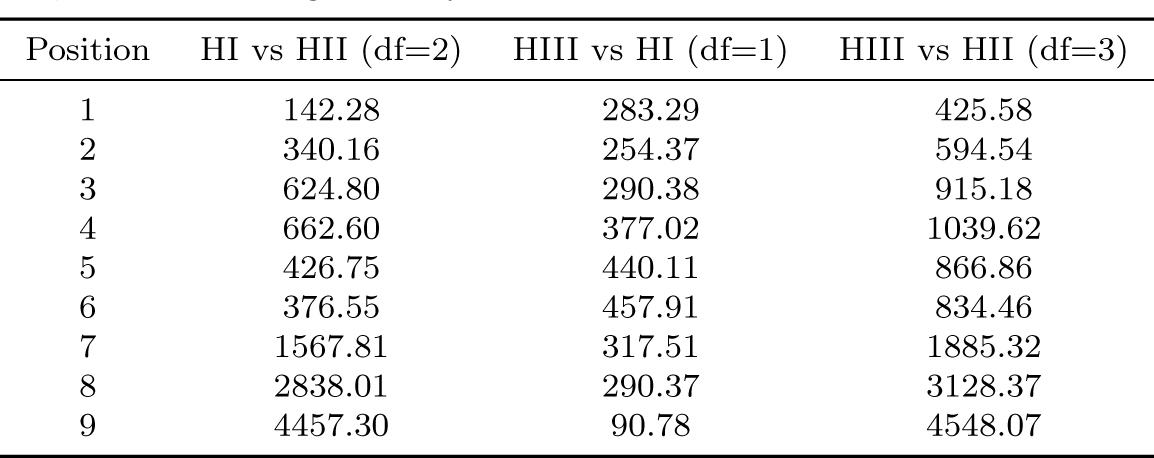
Likelihood ratio test statistic (LRT= -2*ln(likelihood ratio)) for different models fit to autosomal data. Differences in number of parameters are given as the degrees of freedom in brackets. Values are significant for all positions and comparisons. Both HI and HIII fit significantly better than HII, and HIII fits significantly better than HII.

| Position | HI vs HII (df=2) | HIII vs HI (df=1) | HIII vs HII (df=3) |
| --- | --- | --- | --- |
| 1 | 142.28 | 283.29 | 425.58 |
| 2 | 340.16 | 254.37 | 594.54 |
| 3 | 624.80 | 290.38 | 915.18 |
| 4 | 662.60 | 377.02 | 1039.62 |
| 5 | 426.75 | 440.11 | 866.86 |
| 6 | 376.55 | 457.91 | 834.46 |
| 7 | 1567.81 | 317.51 | 1885.32 |
| 8 | 2838.01 | 290.37 | 3128.37 |
| 9 | 4457.30 | 90.78 | 4548.07 |

**Table S3:** Likelihood ratio test statistic (LRT= -2*ln(likelihood ratio)) for different models fit to X chromosomal data. Differences in number of parameters are given as the degrees of freedom in brackets. Values are significant for all positions and comparisons. Both HI and HIII fit significantly better than HII, and HIII fits significantly better than HII.

| Position | HI vs HII (df=2) | HIII vs HI (df=1) | HIII vs HII (df=3) |
| --- | --- | --- | --- |
| 1 | 43.55 | 52.84 | 96.39 |
| 2 | 34.06 | 46.09 | 80.15 |
| 3 | 55.98 | 67.98 | 123.95 |
| 4 | 82.40 | 60.82 | 143.22 |
| 5 | 65.87 | 67.41 | 133.28 |
| 6 | 39.52 | 73.20 | 112.72 |
| 7 | 228.29 | 44.89 | 273.18 |
| 8 | 260.64 | 44.90 | 305.55 |
| 9 | 525.97 | 21.12 | 547.10 |

